# A postnatal human lung developmental atlas reveals windows of genetic vulnerability to chronic lung disease

**DOI:** 10.64898/2026.03.20.713166

**Authors:** Joseph D. Planer, Michael P. Morley, Songlin Cai, Andrew R. Peterson, Minqi Lu, Yifan Yang, Hannah Hallquist, Sarah E. Schaefer, Justine Carl, Su Zhou, Susan M. Lin, Prashant Chandrasekaran, Lisa R. Young, Edward Cantu, William Peranteau, David B. Frank, Edward E. Morrisey, Maria C. Basil

**Affiliations:** Department of Medicine, Perelman School of Medicine, University of Pennsylvania, Philadelphia, PA 19104, USA; Penn-Children’s Hospital of Philadelphia Lung Biology Institute, University of Pennsylvania, Philadelphia, PA 19104, USA; Penn Cardiovascular Institute, University of Pennsylvania, Philadelphia, PA 19104, USA; Department of Cell and Developmental Biology, University of Pennsylvania, Philadelphia, PA 19104, USA; Division of Pediatric Cardiology, Department of Pediatrics, Children’s Hospital of Philadelphia, University of Pennsylvania, Philadelphia, PA 19104, USA; Division of Pulmonary and Critical Care Medicine, Department of Medicine, Washington University in St. Louis School of Medicine, University of Pennsylvania, Philadelphia, PA 19104, USA; Division of Pulmonary and Sleep Medicine, Department of Pediatrics, Children’s Hospital of Philadelphia, University of Pennsylvania, Philadelphia, PA 19104, USA; Department of Surgery, University of Kentucky

**Author notes:** Corresponding author: Maria C. Basil, M.D., Ph.D., University of Pennsylvania, Translational Research Center, Room 11-112, 3400 Civic Center Boulevard, Building 421, Philadelphia, PA 19104-5129.

## Abstract

At birth, the lungs undergo an abrupt physiologic change, as the function of gas exchange transitions from the placenta to the lung. Subsequent postnatal development of the lungs is marked by a rapid and profound increase in the growth of the distal airways and alveolar gas exchange compartment. Insults during this period increase the risk of developing lung disease later in life, though how early-life events affect adult disease onset remains unclear. We generated a single-cell atlas of postnatal human lung development from birth through adulthood and mapped temporally regulated gene expression changes in each cell lineage. Using this atlas, we identified disease risk-associated genes with developmentally regulated expression. These analyses reveal cell type-specific and temporally restricted expression of genes associated with adult lung disease risk, including COPD. Heritability enrichment analysis demonstrated that COPD genetic risk is enriched in genes active during early postnatal endothelial development, linking early-life vascular maturation to adult disease susceptibility. These findings characterize the early window of susceptibility for adult chronic lung diseases and establish a framework to guide mechanistic studies of disease-associated genes.

## Main Text

Respiratory insults during early life, from prematurity to poor air quality to respiratory infection, are associated with a substantial increase in the risk of developing chronic obstructive pulmonary disease and other chronic lung diseases in adulthood^1–4^. Lower respiratory tract infections before the age of two nearly double the risk of premature death from respiratory disease, while childhood pneumonia is associated with a 40% increased risk of adult COPD^5^, reduced lung function, and radiographic evidence of airway disease decades later^6^. Despite this well-established link, the molecular mechanisms that regulate postnatal lung growth and render the developing lung vulnerable to injuries that contribute to chronic lung disease remain poorly understood.

This period of vulnerability coincides with a dramatic phase of human lung growth and maturation. At birth, the function of gas exchange is abruptly transitioned from the placenta to the respiratory system as the lungs fill with air for the first time. This transition is followed by rapid development and growth of the lung over the next months and years, resulting in an approximately twenty-fold increase in the lung’s surface area^7,8^. The number of respiratory airways and alveoli increases and matures to support the essential gas exchange function of the lungs, which increases throughout childhood to match the metabolic demands of the growing body. Given the significant cellular heterogeneity in the developing and mature respiratory system, postnatal maturation likely requires coordinated regulation of epithelial, endothelial, mesenchymal, and immune development.

After acute injuries the lung undergoes a significant regenerative response, as most insults and illnesses resolve with the restoration of normal lung function^9,10^. In contrast, once pulmonary function is lost in chronic lung disease, it is generally not reacquired with current therapies, which are not designed to target regeneration^11^. Since the postnatal period of lung development and growth has an important impact on long-term respiratory function and disease susceptibility, it is imperative to understand the cellular and molecular pathways that regulate normal development. This knowledge will provide a framework for identifying how cellular and molecular responses can be harnessed to promote regeneration in the human lung.

To identify the cellular and molecular events defining human postnatal lung growth and development, we generated a transcriptional human lung cell atlas that spans the day of birth through adulthood. These data, along with an accompanying web visualization, constitute a novel resource for the developmental biology and pulmonary research communities. We demonstrate that multiple transcriptional pathways are differentially regulated across postnatal life in various lineages. Critically, we identify genes associated with adult chronic lung disease risk through genome-wide association studies (GWAS) that exhibit cell type- or lineage specificity and are temporally restricted during postnatal development. To determine whether disease-associated genetic variants are enriched in developmentally regulated genes, we applied partitioned heritability analysis to gene sets defined by their temporal and lineage-specific expression patterns^12^. We identify specific cellular contexts and developmental time points that represent periods of heightened vulnerability, offering new targets for prevention strategies that could intervene during these critical windows to reduce the lifelong burden of chronic lung disease. This analysis revealed enrichment in early-expressed endothelial genes for COPD risk. This molecular framework suggests that a subset of genetic risk factors that predispose individuals to adult lung disease act during early life, potentially affecting development or increasing susceptibility to respiratory insults that permanently alter lung structure and function.

### Construction of The Human Postnatal Lung Development Atlas

To generate an atlas of human postnatal lung development, we collected pediatric lung tissue samples through the Penn-CHOP Lung Biology Institute Human Lung Tissue Biobank. We categorized samples into six developmental stages: Day Zero Neonate (Day 0), Neonate, Infant, Toddler, Teen, and Adult. A subset of these samples also has accompanying proximal airways or separate sequencing of the CD45^+^ fraction (see Methods for additional details) (**Fig. 1S1a,b**). In total, 313,150 cells from 30 individuals were analyzed by scRNAseq. Cellular lineages were annotated based on expression of lineage-specific canonical marker genes in cell clusters (**Fig. 1S2a-d, Fig. 1S3a**). Next, the major lineages were subset and reannotated based on the expression of cell type-specific canonical marker genes (**Fig. 1S2e, 1S3b**). To ensure our annotation was consistent with established large-scale data sets, we separately performed reference-based annotation using both the Human Lung Cell Atlas (HLCA)^13^ and LungMap Single Cell Reference (LMCR)^14^. There was generally high concordance between the three annotation schemes for all lineages, with the endothelium and mesenchyme showing the greatest divergence (**Fig. 1S4**). All expected lung cell types were present along with a small group of Schwann cells expressing *MPZ*, and chondrocytes expressing *COL2A1* (**Fig. 1S2e, 1S3b**)^15,16^. Certain cell types were present at some time points and not others, such as alveolar macrophages (aMACs), which are absent at birth and accumulated during the first weeks to months of life, as has been reported^13,17,18^ (**Fig. 1S3c, 1S5**).

### Patterns of age-associated transcriptional change

We generated pseudobulk expression profiles for each cell type in each sample and used a non-linear model to define age-associated differentially expressed genes (DEGs)^19,20^. After clustering, we observed multiple patterns of temporal gene expression change, including genes that are down-regulated or up-regulated during the first days of postnatal life or later in childhood, along with genes whose expression changed later during development (**Fig 1a**).

**Figure 1:**
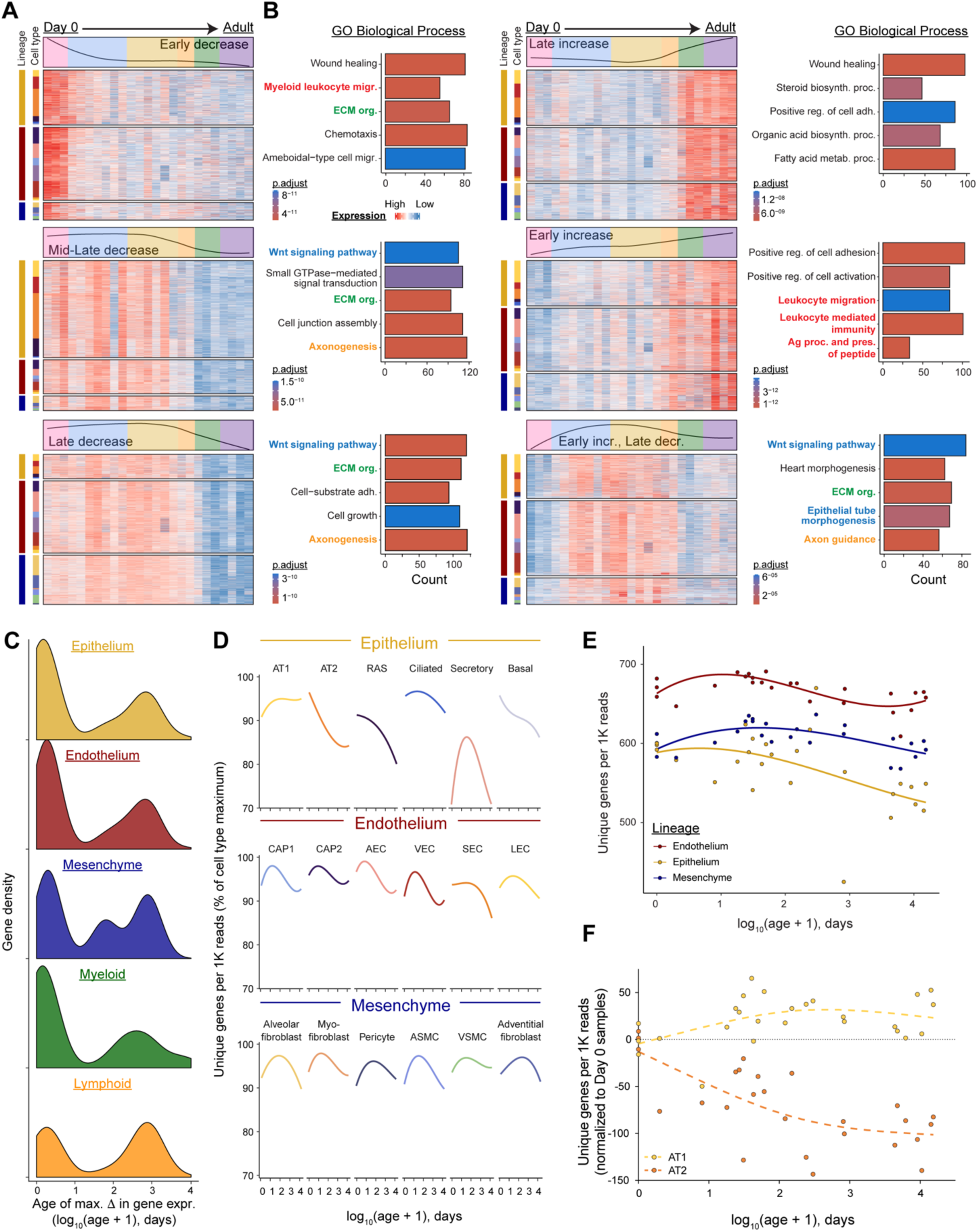
Patterns of gene expression in a human pediatric developmental transcriptional atlas. (**a**) Differentially expressed genes (DEGs) for each cell type were defined using a spline fit model and grouped into 6 clusters using k means. Samples are arranged from left to right in order of increasing chronologic age with a rolling window of relative gene expression level in the top annotation. Color segments in the top annotation box correspond to developmental stages according to the color scheme used in Fig. 1**S****3c**. Left annotation bars denote lineage and cell type, with color schemes corresponding to Fig. 1**S****3a,b**. (**b**) GO:Biologic Process over-enriched categories within each cluster. Text is colored according to theme: red, inflammation and immune signaling; blue, Wnt and epithelial growth signaling; green, extracellular matrix organization; gold, axon and neuron development. (**c**) Density histogram for DEGs plotted by the age of maximal change in transcriptional abundance. (**d**) The number of unique genes per 1000 was determined for each cell. Curves were generated for each cell type after normalizing for the maximum value per cell type. (**e**) Median values for each lineage and donor are plotted and colored according to lineage. (**f**) The median number of unique genes per 1000 in AT1 and AT2 cells from each donor are shown over age. Cell types in (a) are color coded according to the schematic in panel (d).

To define which transcriptional programs were changing over development, we performed Gene Ontology (GO) overenrichment analysis on each cluster. We found that changes in Wnt signaling, epithelial growth, and extracellular matrix (ECM) synthesis were coordinated across several cell types and downregulated in phases as the lung matured (**Fig. 1b**), correlating with previously observed changes in mouse lung development. Multiple phases of change in immune signaling gene expression were evident, with genes involved in leukocyte migration were downregulated after peaking in Day 0 neonates while genes involved in adaptive immune signaling were upregulated in teens and adults (**Fig. 1a,b**).

To quantify when cellular behavior was changing most dramatically during development, we defined the time point of maximal transcriptional change for each DEG in each cell type or lineage. Lineage-level analysis revealed two main windows of temporal change: the first occurring from day 0 (birth) through the first month, and a second at the transition from childhood to adolescence. These periods of change were consistent across different cell types and were also present when we compared across developmental stages, suggesting a coordinated transcriptional response at different points in postnatal development (**Fig. 1c, 1S6a,b**). These time points correspond to two major physiological shifts: the transition from placental to pulmonary oxygenation and the resolution of the growth phase of the respiratory system.

### Cellular transcriptomic diversity is age- and lineage-dependent

Prior work has shown that transcriptional diversity is a surrogate for the complexity of cellular behavior^21,22^. We hypothesized that the cells of the lung would demonstrate less functional complexity, corresponding to a decrease in transcriptomic diversity, as the lung adopted its adult form and function.

To test this hypothesis, alpha diversity metrics were calculated for each cell type in each sample. We reasoned that cells with a limited number of functions, such as erythrocytes and plasma cells, would exhibit a less diverse transcriptome than cells with a larger number of known functions. Indeed, this was reflected in our analysis of adult lungs from our data set (**Fig. 1S7a,b**). Among the structural (non-immune) lineages, the epithelium showed the greatest transcriptomic diversity (**Fig. 1S7a,b)**.

Analysis of age-associated patterns revealed several distinct trajectories of cellular transcriptomic diversity. Within the endothelium and mesenchyme, cell types typically shared a similar temporal pattern (**Fig 1d)**. Epithelial cell types exhibited a broad range of temporal patterns (**Fig. 1d,e, 1S7a**). Within the alveolar epithelial compartment, AT1 transcriptomic richness stayed high or increased slightly with age (**Fig. 1f**). This contrasts with the progenitor populations of the epithelium, with alveolar type 2 (AT2), respiratory airway secretory (RAS), and basal cells all exhibiting a gradual reduction in the number of unique genes detected over time (**Fig. 1d, 1S6a**). Together, these data support a dynamic set of transcriptional changes regulated both at the lineage and cell-type-specific levels, demonstrating the multifaceted processes governing postnatal lung growth and maturation.

### Postnatal changes in alveolar epithelium associate with ECM generation and remodeling

The human respiratory epithelium comprises two morphologically and functionally distinct alveolar epithelial cell types, AT1 and AT2 cells, as well as respiratory airway secretory (RAS) cells, which line the distal airways^23–25^. To identify the major developmental transcriptional changes in the respiratory epithelium, we generated scores for cellular signaling pathways in the Molecular Signatures and PROGENy databases and defined the subset that changed over development (**Fig. 2S1**). This analysis demonstrated downregulation of TGF-β signaling in the respiratory epithelium, and upregulation of genes involved in xenobiotic and fatty acid metabolism over development. We paired this analysis with gene set enrichment analysis, which also identified changes in canonical developmental pathways such as Wnt ^26–28^ and TGF-β signaling in the respiratory epithelium as previously reported in the mouse lung^29^ (**Fig. 2S2a-f**). We also observed that ECM organization was upregulated at early developmental time points across the entire distal epithelium (**Fig. 2S2a-c**).

Since the balance of TGF-β signaling coordinates cell fate decisions in the distal airways and alveoli^27–30^, we interrogated this fundamental pathway in our dataset. We generated a TGF-β activity score using downstream transcriptional targets along with an activity score for SMAD3, a transcription factor that transduces TGF-β signals. Within the respiratory epithelium, TGF-β signaling was strongest in AT1 and RAS cells, consistent with its described importance in AT1 function and maintenance (**Fig. 2a,b**)^29,30^. Both TGF-β and SMAD3 activity demonstrated a similar trend, consistent with downregulation of TGF-β activity in the distal epithelium through development (**Fig. 2c**).

**Figure 2:**
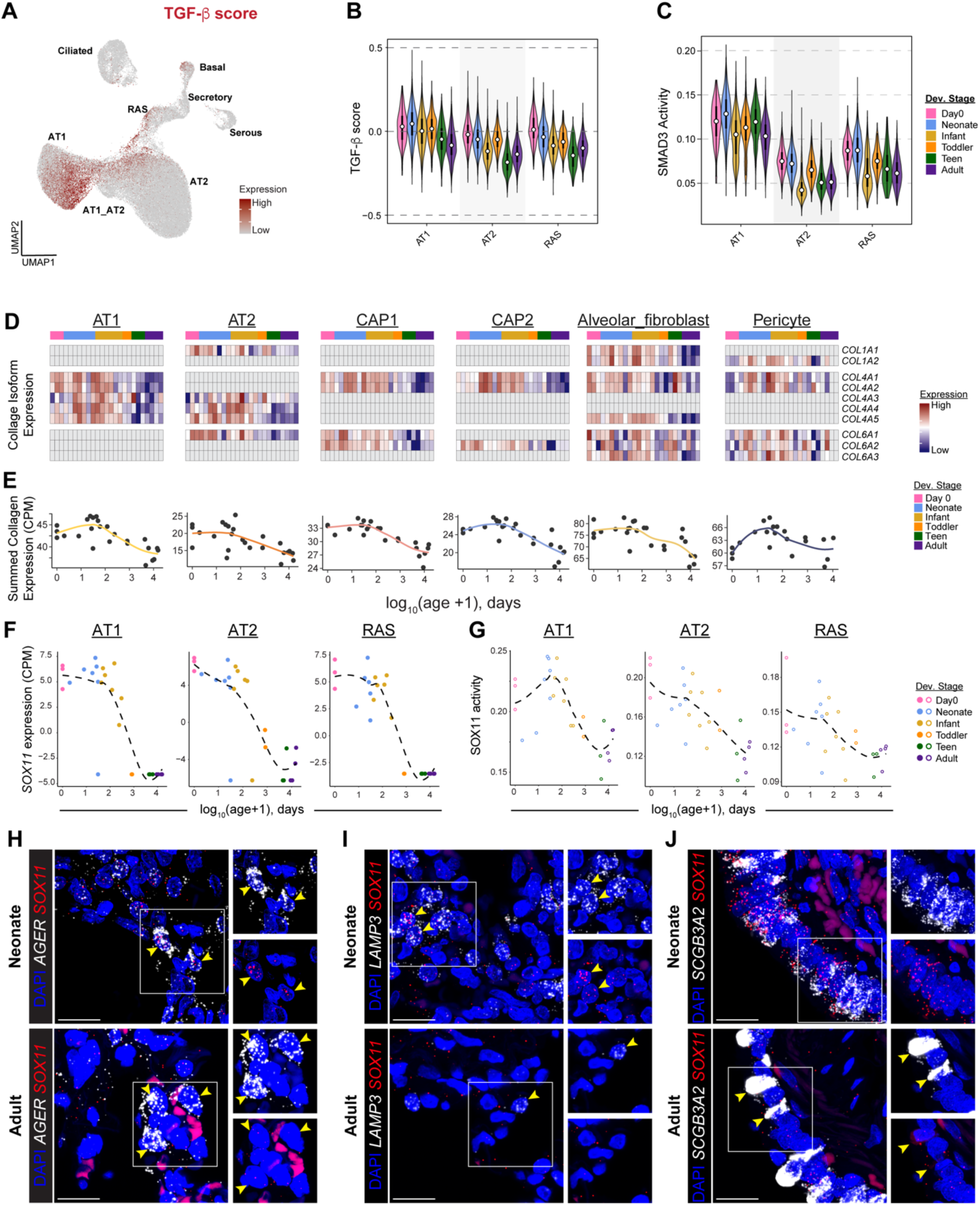
Postnatal regulation of epithelial developmental signaling pathways. Genes expressed downstream of TGF-β were used to generate a TGF-β score for each cell in the original parent set. This score was plotted (**a**) and summarized by age for the three principal respiratory epithelial cell types (**b**). (**c**) SMAD3 activity was calculated and summarized as in (b). (**d**) Expression of individual collagen I/IV/VI genes across major alveolar cell types. (**e**) Summed collagen expression for these cell types. *SOX11* expression (**f**) and activity scores (**g**) were plotted for each cell type over postnatal development. (**h-j**) RNAscope imaging of *SOX11* and cell type specific markers for neonates and adults.

TGF-β has a well-established role in promoting ECM synthesis and remodeling^29,31,32^. The orderly deposition of ECM is critical to lung maturation and growth, and ECM abnormalities are a feature of many chronic lung diseases, such as emphysema and idiopathic pulmonary fibrosis (IPF)^33,34^. We observed that transcription in the collagen family of genes was downregulated with age across several cell types and lineages in the alveolar niche as has been observed in proteomic data (**Fig. 2d**). To further interrogate ECM production and signaling, we generated a composite score of genes in collagen families I/IV/VI, which demonstrated that collagen transcript abundance peaks during the first 100 days of life (**Fig. 2e**). These data suggest that ECM establishment is a complex, multicellular and multilineage process that increases during phases of rapid lung growth and declines when growth slows. This temporal pattern parallels TGF-β pathway activity, supporting a coordinated role for TGF-β signaling in regulating ECM deposition during postnatal lung development.

### Transcriptional targets of TGF-β and Wnt signaling are temporally regulated during postnatal lung development

Intriguingly, the SOX C family transcription factor *SOX11*, was downregulated with age across multiple lineages (**Fig. 2f, 2S2d-i**). SOX11 is required for branching morphogenesis during prenatal lung development^35,36^ and is a downstream target of Wnt and TGF-β signaling^37–39^. TF activity scores corroborated that SOX11 activity mirrors *SOX11* expression in these cell types during postnatal development (**Fig. 2g**). This decrease in *SOX11* expression was confirmed using RNAscope analysis (**Fig. 2h-j**). These findings suggest that SOX11 may play a role in regulating a shared transcriptional program within the epithelium during postnatal lung development.

The shared transcriptional programs across the early lung distal epithelium raised the question of cellular specification and whether cells becoming more fixed in their final state as they mature. We noted that the respiratory epithelial cell populations progressively separated from each other in UMAP space over development (**Fig. 1S5b**). Re-clustering AT2 and RAS cells revealed a transcriptionally intermediate state enriched at the time of birth (**Fig. 2S3a,b**). This RAS-AT2 intermediate state, which is similar to states observed in chronic lung disesase^23,40^, was greatly reduced or lost shortly after birth (**Fig. 2S3b**). Pseudotime trajectory analysis demonstrated the downregulation of *SOX4* and upregulation of *ETV5* along the RAS-AT2 differentiation trajectory as has been described^23^, while the aryl hydrocarbon receptor (*AHR*) was upregulated during the transition from RAS to AT2 (**Fig. 2S3c,d**). We similarly observed that AT2 and AT1 cells were much less transcriptionally distinct during the first weeks of postnatal life compared to older time points (**Fig. 2S3e-g**). These data suggest that as the lung matures during development, the cell populations of the distal lung become more distinct with less shared cellular and transcriptional programs.

### Coordinated development of the alveolar endothelial lineage

The alveolar niche comprises a coordinated network of epithelial and endothelial cells that develop in parallel to establish the gas exchange interface. To define how cellular crosstalk regulates vascular development, we performed ligand-receptor analysis on cells within the alveolar niche, focusing on pathways known to regulate angiogenesis^41–45^ (**Fig. 3S1a,b**). AT1 cells emerged as a central regulator of endothelial development through multiple signaling axes (**Fig. 3S1c**). AT1 cells represented the major cellular source of *VEGFA* transcript^29,46^, which decreased over time, while CAP1 and CAP2 cells expressed the VEGF receptors *KDR* and *FLT1* at similar levels across developmental age (**Fig. 3S1d**)^47,48^. This pattern suggests that angiogenic activity during development is regulated in part by AT1 cells, which dynamically regulate ligand expression rather than by endothelial populations modulating endothelial receptor availability.

Additional cell-cell signaling pathway analysis identified other pathways that have been previously shown to regulate angiogenesis (**Fig 3a-d**). Notch signaling plays a central role in sprouting angiogenesis, with DLL4 transducing VEGF-A signaling to promote orderly branching^45,49^. Notch pathway signals were enriched in CAP1 cells, with expression of DLL4 decreasing over age (**Fig. 3a**). Semaphorin-plexin-neuropilin signaling plays a central role in regulating endothelial vessel branching and pruning^41^. The *PLXND1* receptor, expressed by CAP1 cells, was significantly downregulated with age (**Fig. 3b**). Similarly, expression of midkine^50^ and endothelin in CAP1 cells decreased with maturation (**Fig. 3c,d**). Together, these findings suggest that vascular development is coordinated through multiple, temporally synchronized signaling pathways, with AT1 and endothelial cells serving as key regulatory hubs during postnatal angiogenesis.

**Figure 3:**
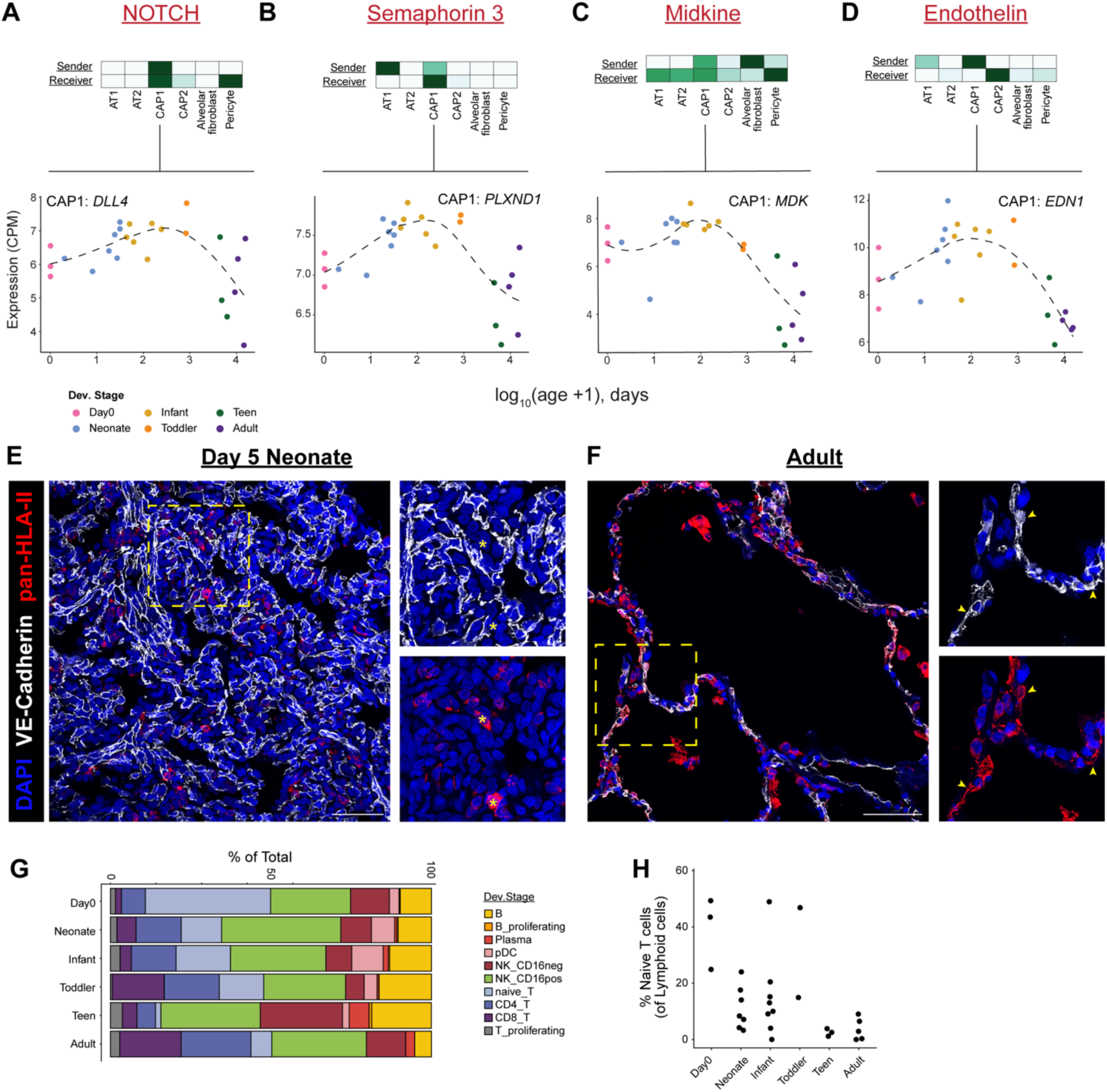
Vascular patterning and immune-endothelial signaling through development. Pathway signaling flow and major ligand expression patterns in Notch (**a**), Semaphorin 3 (**b**), Midkine (**c**), and Endothelin (**d**) pathways. (**e,f**) IF imaging of VE-Cadherin and HLA-II in neonate and adult samples. (**g,h**) Proportional representation of lymphocyte subsets over developmental stages.

### Endothelial inflammatory signaling peaks during both the perinatal period and the transition to adulthood

Beyond establishing the capillary network for gas exchange, we observed that the endothelium also exhibited dynamic changes in immune signaling throughout development with two peaks of transcriptional variation: 1) the first days of postnatal life and 2) in late development as the lung stop growing in size (**Fig. 1c**, **Fig. 1S6**). The first phase was characterized by increased expression of genes associated with inflammation (including responses to lipopolysaccharide (LPS), wounding, and angiogenesis) in day 0 samples (**Fig. 3S2a,b**). Upregulation of cytokines involved in innate immune signaling (*CCL2, CXCL1, IL6, CSF3*), markers of cell activation (*NFKB1, RELA*), and cellular stress (*HSF1, XPB1*) were also observed at the time of birth (**Fig. 3S2a,b,d**). This is consistent with major cellular stress in the vasculature of day 0 neonates, as the pulmonary endothelium assumes the role of gas exchange and experiences the stress of parturition.

The second phase of endothelial transcriptional change was characterized by an upregulation of genes involved in adaptive immune responses. Endothelial cells have been observed to express major histocompatibility complex II (MHC-II), although its functional significance remains unclear^51–53^. As the lung reaches maturity, multiple genes in the human leukocyte antigen class II (HLA-II) locus, which encodes human MHC-II, were upregulated in the endothelium (**Fig. 3S2a,c,e**). IF imaging of HLA-II protein demonstrated a paucity of endothelial MHC-II expression in neonates, in contrast to adults where robust MHC-II colocalization with VE-Cadherin was evident (**Fig. 3e,f**).

Given this finding, we hypothesized that these changes may be associated with the maturation of the adaptive immune system within the lung. To test this, we further refined our taxonomic classification of lymphoid immune cells in the lung (**Fig. 3S3a,b**). We observed that the percentage of naïve T cells declined with age, while the percentage of mature and memory T cells, which are nearly absent at birth, increased with time (**Fig. 3g,h**). This suggests that maturation of lymphocyte populations in the lung occurs in parallel with upregulation of EC-specific MHC-II expression, which may have important implications in lung diseases characterized by chronic immune dysregulation^54–56^.

### Conservation of the imprinted gene network between species links postnatal growth to disease susceptibility

The developmental programs described above, including coordinated regulation of TGF-β signaling, ECM deposition, and cellular specification in the alveolar niche, define a period of rapid postnatal growth during which the lung undergoes profound structural and functional change. Early life insults during this period, from prematurity to respiratory infection, are associated with substantially increased risk of chronic lung disease in adulthood. However, mechanistically linking specific developmental programs to later disease risk requires identifying which molecular mechanisms are conserved across species and could therefore be perturbed in model systems.

We hypothesized that, despite significant differences in murine and human lung physiology and anatomy, core developmental programs would be conserved across species. To test this, we leveraged two existing scRNA-seq datasets that characterized postnatal lung development in mice (**Fig 4S1**)^57,58^. We reasoned that understanding the molecular regulators governing the tempo of postnatal lung growth may reveal how disruption of these programs during critical developmental windows leads to permanent alterations in lung structure and function. To identify these regulating factors, we focused our analysis on cell types within the alveolar niche that have facultative progenitor capacity: AT2s, CAP1s, and alveolar fibroblasts^46,59,60^. For each cell type, we defined the subset of genes that were significantly up- or downregulated across all three developmental datasets, identifying a conserved transcriptional response during postnatal lung development (**Fig. 4a,c,e**).

**Figure 4:**
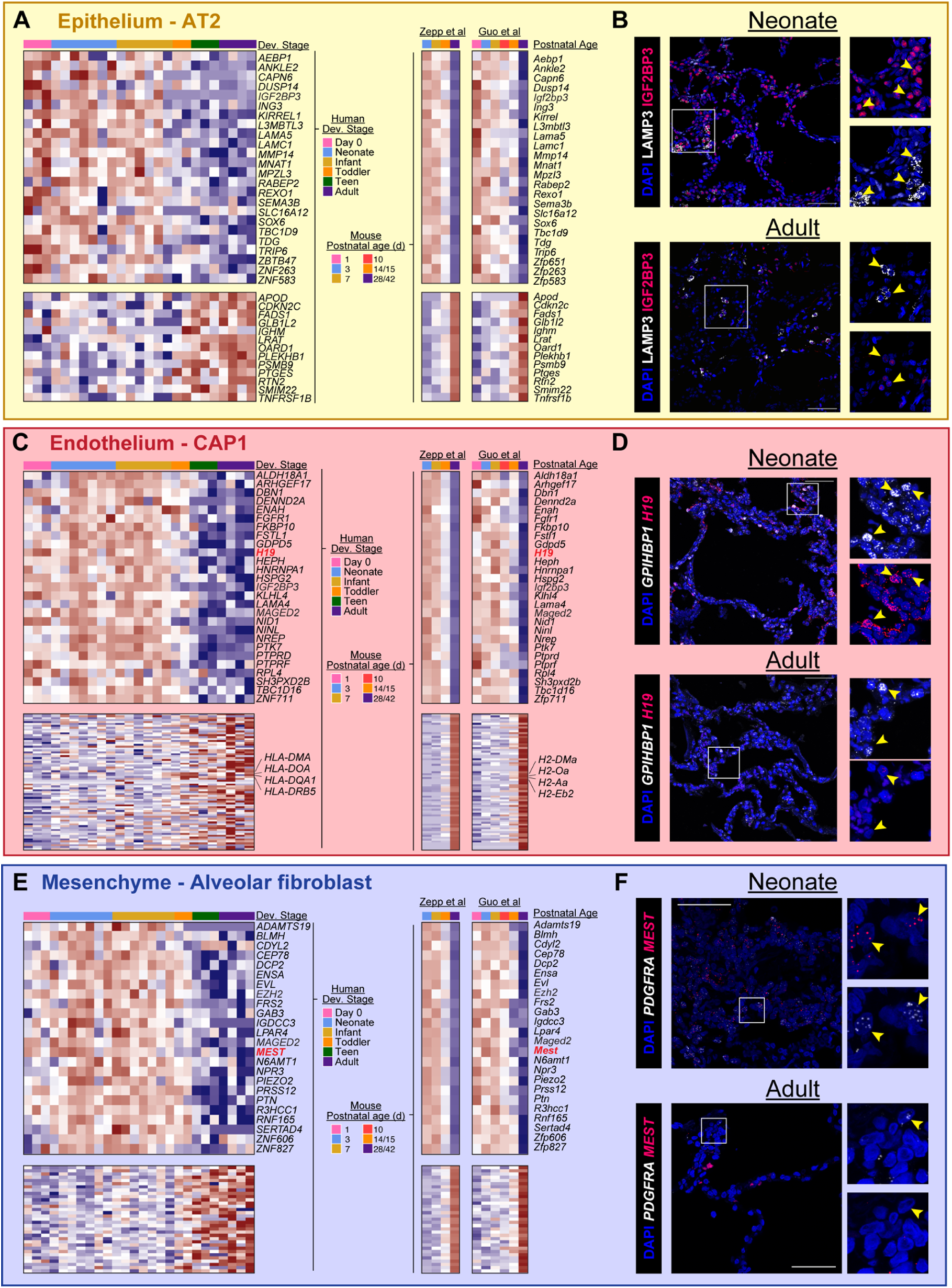
Conserved patterns of age-associated gene expression between humans and mice. Shared patterns of age-associated expression were defined for AT2 cells, CAP1 cells, and alveolar fibroblasts. Genes that were significantly downregulated during postnatal development across all three data sets are plotted (**a,c,e**). IF imaging of IGF2BP3 in AT2 cells (**b**), *H19* in CAP1 cells (**d**), and *MEST* in alveolar fibroblasts (**f**) across age in humans and mice. Genes highlighted in red are components of the IGN.

A growing body of evidence has identified a coordinated imprinted gene network (IGN) that coordinates pre- and postnatal somatic growth across multiple organs. Expression of canonical IGN members. including *Igf2*, *H19*, *Mest*, *Peg3*, *Dlk1*, and *Cdkn1c*, declines coordinately during growth deceleration in mouse lung, kidney, and heart, with temporal patterns that parallel the decline in organ growth rate^61–64^. This network can also be pathologically reactivated: loss of imprinting (LOI) at the *IGF2* locus has been proposed to drive autocrine signaling in AT2 cells during chronic injury, producing an imbalance in proliferation and differentiation that results in emphysema or malignant transformation^65,66^. However, despite evidence that the IGN is active in the postnatal lung, no study has resolved expression patterns of network members during human lung development, a gap that limits our ability to connect specific developmental programs to cell-type-specific disease mechanisms.

Among the conserved developmentally downregulated genes, we identified multiple canonical components of the IGN distributed across distinct cell types of the alveolar niche (**Fig. 4, 4S2**). Of the core group of 11 genes initially reported as part of the IGN^67^, several were downregulated across many cell types (e.g. *H19*, *MEG3*), while others exhibited greater cell-type specificity (*DLK1*, *SLC38A4*; **Fig. 4S2**). In CAP1 cells, we observed conserved downregulation of *H19*, a maternally expressed long non-coding RNA and one of the most well-characterized IGN members (**Fig. 4c**). RNAscope imaging confirmed higher *H19* transcript levels in CAP1 cells of neonates compared to adults in both humans and mice (**Fig. 4d**). In alveolar fibroblasts, we observed a striking downregulation of *MEST* (formerly PEG1), a paternally expressed imprinted gene whose postnatal decline has been observed in multiple murine tissues^61^ (**Fig. 4e,f**). Alongside these canonical IGN members, we observed the conserved developmental downregulation of IGF2BP3 in both AT2 and CAP1 cells (**Fig. 4a,c**). IGF2BP3 is a member of the IGF2 mRNA-binding protein family that post-transcriptionally regulates the stability and translation of a broad repertoire of growth-associated transcripts, including but not limited to *IGF2*^68^. IF imaging of IGF2BP3 demonstrated strong staining in multiple cell types in neonates, including AT2 cells, while staining in adult samples was substantially reduced (**Fig. 4b**). Notably, *IGF2* transcript itself was not detected at appreciable levels in AT2 cells in our datasets, suggesting that the developmental role of IGF2BP3 in this compartment likely operates through regulation of other target mRNAs rather than through canonical IGF2 signaling. The coordinate decline of IGF2BP3 alongside canonical IGN members nevertheless raises the possibility that its expression is subject to shared upstream regulatory inputs during postnatal growth deceleration.

The coordinated decline of multiple growth promoting pathways suggests that postnatal growth deceleration in the lung involves a broad transition in growth capacity rather than the modulation of a single signaling axis. This developmental window, during which multiple growth-promoting programs are simultaneously active, may represent a period of heightened vulnerability to environmental insults that could permanently alter the trajectory of alveolar maturation. The IGN may also intersect with other developmental programs identified in this study. PLAGL1, a transcriptional regulator within the network, has been shown to directly regulate ECM genes including collagens, providing a potential link between the IGN and the coordinated ECM remodeling we observed in the developing alveolar niche^69^ (**Fig. 2d,e**). The conservation of these programs across species validates the use of murine models for mechanistic investigation, while the cell-type resolution provided by our atlas identifies the specific cellular contexts in which perturbation studies can be performed.

### Defining the Genetic Predisposition for Chronic Lung Disease across the lifespan

Chronic lung diseases have complex genetic and environmental etiologies, with heritability estimates of ∼50% for COPD and ∼20% for lung cancer^70–72^. Genome-wide association studies (GWAS) have identified over 80 loci associated with COPD risk alone^73^; however, determining the mechanistic role of these variants has been challenging due to limitations of animal models. Several studies have linked COPD susceptibility to reduced lung function in early life independent of exposure^74,75^, suggesting that a component of genetic risk for adult chronic lung disease may be established during development. Given the plethora of divergent risk factors for diseases like COPD, endotyping these complex diseases is essential. Asthma, a chronic lung disease with well-validated endotypes^76,77^, has seen great success with therapies that are targeted to specific pathologic mechanisms in defined patient populations. Replicating this success in COPD and other chronic lung diseases has proven more elusive.

Having defined the cellular and molecular landscape of human postnatal lung development, we next investigated whether these cellular and developmental programs harbor genetic risk variants for adult chronic lung disease. To investigate how early lung growth might influence the development of chronic lung disease in adulthood, we used the NHGRI-EBI GWAS catalog to generate a list of genes associated with adult chronic lung disease^78,79^. We identified risk-associated genes (RAGs) from the NHGRI-EBI GWAS catalog for multiple chronic lung diseases. For each major cell type and cell lineage, we also identified genes that were significantly down- or upregulated across postnatal development, generating lineage-specific gene sets representing ‘early-expressed’ (subsequently downregulated) and ‘late-expressed’ (subsequently upregulated) programs. These gene lists, which represent cell-type or lineage-specific developmental programs, were compared to RAGs from various lung diseases, allowing us to define age-associated disease-specific RAGs (**Fig. 5a**, **Fig. 5S1**). Many of the RAGs that increase with age were associated with inflammation, including *IL7R* in asthma^80^, *IL1RL1* in asthma and COPD^81,82^, and HLA-II alleles with asthma, COPD, lung cancer, and sarcoidosis^80,83,84^. In contrast, downregulated RAGs were often involved in ECM deposition or processing (*ELN, PRSS35, ADAMTS4*), cellular growth (*SOX11, H19*), or cellular differentiation (*WNT5B*, *FOXP2*)^36,80,85^ (**Fig. 5S1**).

**Figure 5:**
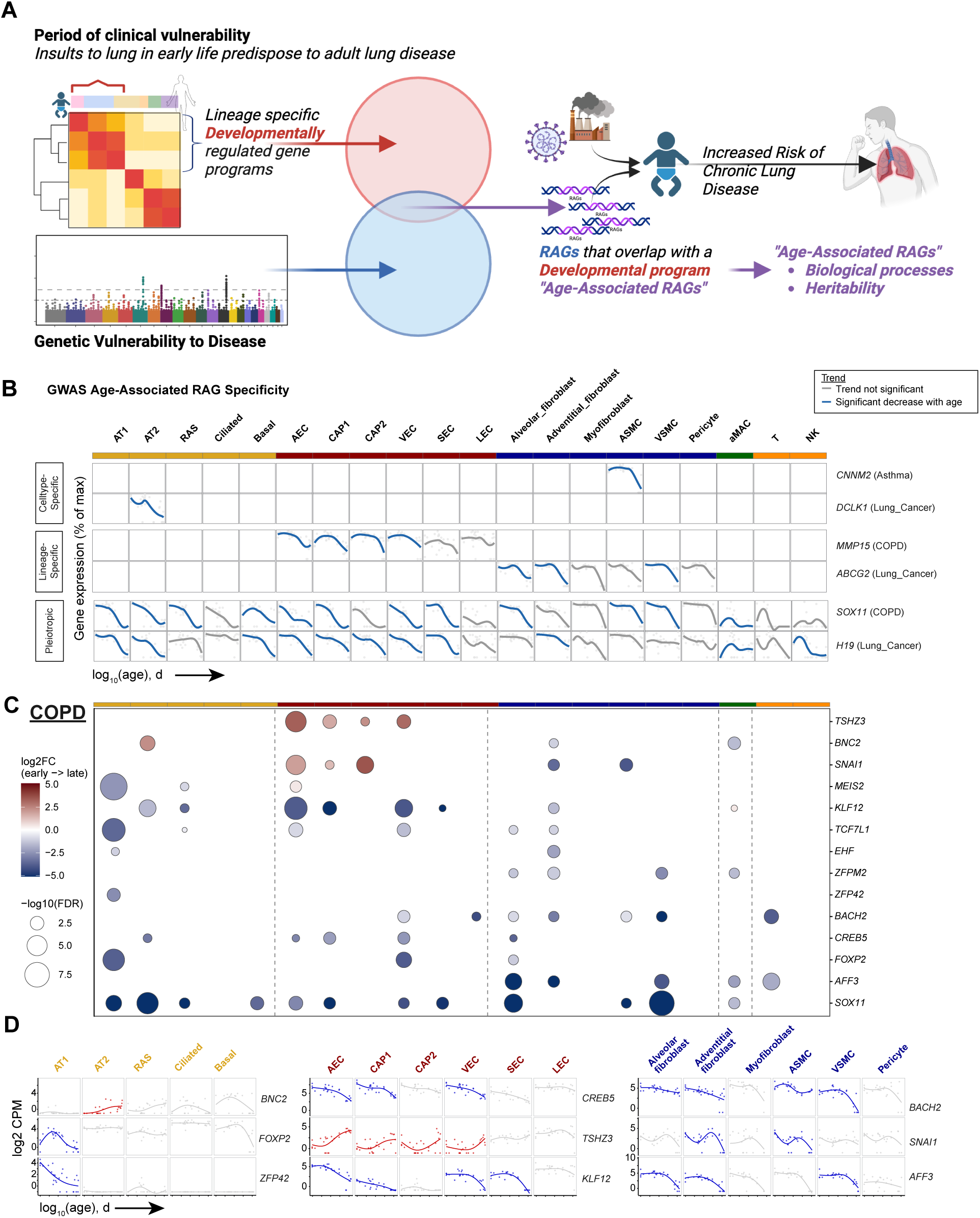
Patterns of age-associated RAG enrichment across lineages. (**a**) Schematic illustrating the strategy for intersecting GWAS RAGs and age-associated DEGs to define disease-specific developmental programs. (**b**) Patterns of specificity in age-associated RAG expression in major lung cell types. (**c**) Age-associated RAG transcription factors in COPD GWAS. (**d**) Patterns of age-associated transcription factor expression corresponding to COPD GWAS hits in select lineages.

We next focused on RAGs associated with early expression and subsequent downregulation, as we reasoned that these transcriptional programs were active when early-life injury and insults can contribute to an individual’s risk for adult lung disease. We further categorized age-associated RAG expression by defining the cell type- or lineage-specificity. 1,354 of age-associated RAGs were cell-type specific, 354 were specific for a single lineage, and 213 were similarly regulated across a broad range of cell types. For example, expression of the magnesium transporter *CNNM2*, which has been associated with age of asthma onset^86^, was significantly downregulated over development in airway smooth muscle cells (**Fig. 5b**). Similarly, *DCLK1* has been associated with stemness in lung adenocarcinoma^87^ and was downregulated over development in AT2 cells, a cell type of origin for adenocarcinoma. Other age-associated RAGs were coordinately regulated across multiple cell types within a lineage, including *MMP15,* which has a well-established association with COPD^88,89^, and ABCG2 across multiple mesenchyme populations (associated with lung adenocarcinoma)^90–92^. In contrast, some early-expressed downregulated RAGs were coordinately regulated across multiple cell types and lineages, such as *SOX11* and *H19* (COPD and lung cancer)^93,94^, suggesting a generalized role in maintaining developmental plasticity or promoting growth during postnatal development (**Fig. 5b**).

We then focused on age-associated transcription factors (TFs) among RAGs as likely central regulators of developmental programs (**Fig. 5c, 5S2**). COPD risk was associated with 14 age-associated TFs, including *ZFP42* (selectively downregulated in AT1 cells), *BNC2* (upregulated in AT2 cells), *CREB5*, and *KLF12* (both downregulated in endothelium) (**Fig. 5c,d)**. Many of these TFs have established roles in lung development, including epithelial specification (*ZFP42*, *BNC2*)^95,96^ and vascular patterning (*CREB5*, *KLF12*)^97,98^, supporting their role in establishing disease vulnerability during critical developmental windows.

We next performed gene ontology over-enrichment analysis of age-associated RAGs by disease. Genes involved in antigen processing and presentation (e.g. MHC-II) were upregulated in endothelium during development and associated with sarcoidosis and asthma (**Fig. 6a**), consistent with our above observations (**Fig. 3**). We further observed that genes involved in epithelial development, which are likely important for the establishment and growth of the alveolus, were strongly associated with lung cancer RAGs (**Fig. 6a**). These included established cancer driver genes such as *DLC1, TP53BP1, ROR1*, and *EZH2*^99–102^ (**Fig. 6b**). In contrast, we observed that endothelial downregulated genes that overlapped with for RAGs for COPD were associated with TGF-β signaling (**Fig. 6a,b**), suggesting these pathways may represent a cell lineage-specific function during development that increases risk for this chronic lung disease.

**Figure 6:**
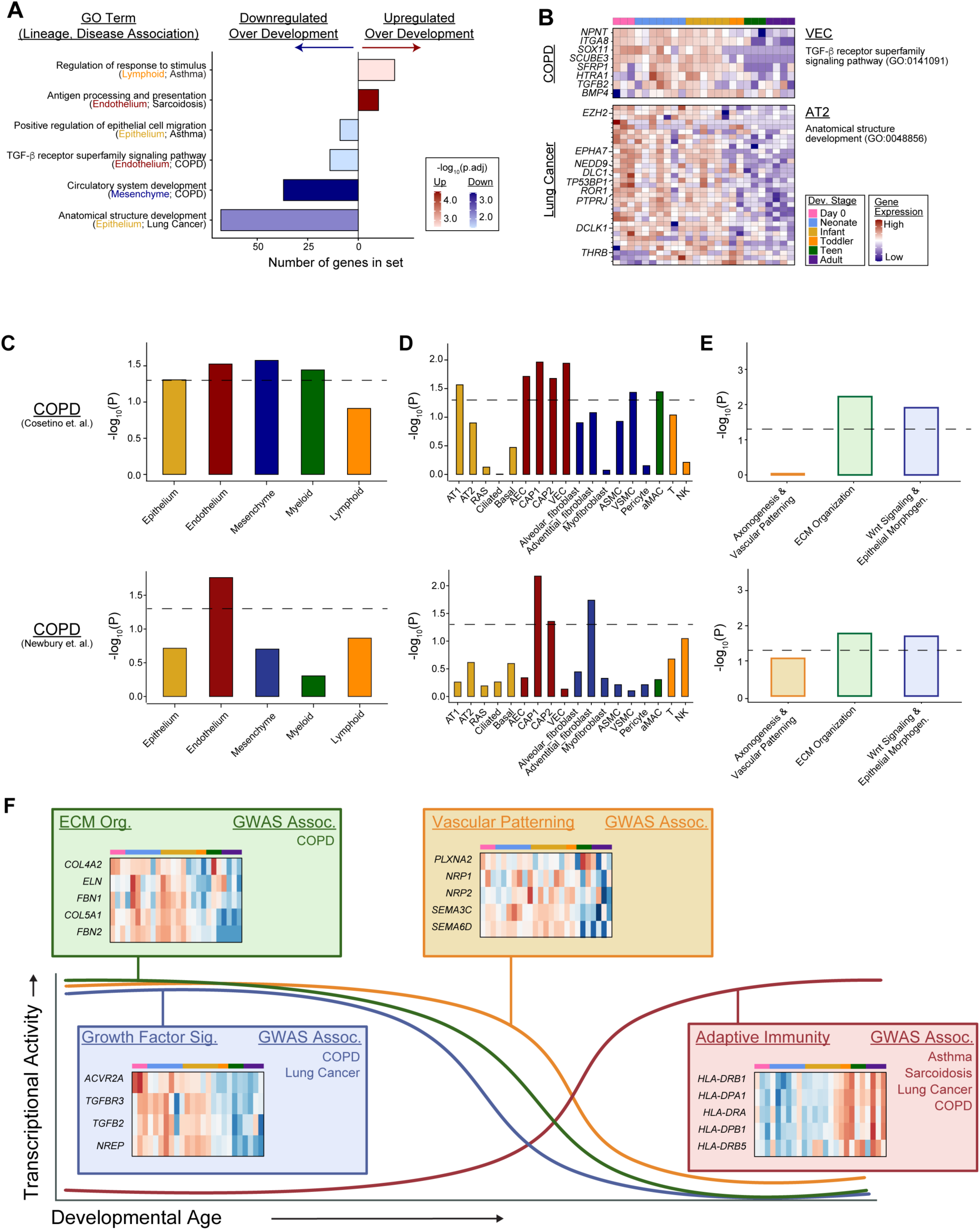
Cell type and lineage-partitioned heritability in asthma and COPD. (**a**) GO:BP over-enrichment analysis of up- and down-regulated gene sets associated with disease bins. (**b**) Expression patterns for genes within enriched GO terms associated with COPD and lung cancer. LDSC regression was performed at the lineage (**c**) and cell type level (**d**) for the early-expressed DEG sets in two separate COPD GWAS studies. (**e**) LDSC regression was performed using gene families derived from GO categories in Fig. 1. Nominal P values are shown with the dashed line denoting P = 0.05. (**f**) Schematic demonstrating age-associated changes in transcriptional activity for major developmental pathways.

### Identifying a molecular and genetic link in endothelial cell programs to COPD

The patterns we observed in age-associated RAGs, including coordinated regulation within specific lineages and developmental windows, suggested that biological processes active during postnatal development might contribute disproportionately to heritable disease risk. To test whether disease-associated genetic variants are enriched in genes with developmentally restricted expression patterns, we applied partitioned heritability analysis^12,103^ to our postnatal lung developmental dataset (**Fig. 6S1**).

We first confirmed whether our dataset could identify cell type- and lineage-specific genetic risk factors for chronic lung diseases, as has been performed previously using the Human Lung Cell Atlas (HLCA)^13^. The HLCA study, which used predominantly European ancestry cohort spanning ages 10-76, identified COPD heritability enrichment in myofibroblasts, as well as asthma heritability in T cells. Using the most mature lung samples from our atlas (teenagers and adults, ages 13-40) we performed partitioned heritability enrichment analysis for each cell type and lineage using linkage disequilibrium (LD) score regression for forced vital capacity (FVC), COPD, and asthma. We began by examining cell-specific heritability using marker gene sets for the most abundant cell types in our data set. Consistent with prior studies, we found significant excess heritability for several mesenchymal cell types associated with FVC, T cells in asthma, and myofibroblasts in COPD (**Fig. 6S2a**).

Given the known associations of early lung injury with the development of COPD, we focused on this disease for our developmental analysis while using asthma as a comparator with known contributions from lymphocyte pathology. We repeated the LDSC analysis in two independent COPD GWAS studies using early- and late-expressed RAGs for each lineage rather than the cell type-specific markers^104,105^. We observed significant heritability enrichment specifically in the early-expressed lymphoid gene set for asthma, consistent with the marker gene analysis (**Fig. S62b-c**). Surprisingly, we also observed a significant heritability enrichment with the early-expressed endothelial gene set, suggesting that a subset of genetic variants that increase COPD risk may be preferentially located near genes active during early postnatal vascular development (**Fig 6c,d)**. Notably, we did not observe any consistent heritability association with late expressed genes, suggesting that these effects are most relevant to genes expressed during active lung development (**Fig. 6S2b,c**). Finally, we wondered if heritability might also be linked to different families of developmentally regulated genes. We generated families of GO:BP terms with similar biological functions and performed LDSC using the subset of age-associated genes from these families. This analysis indicated that genes involved in ECM organization, Wnt signaling, and Epithelial morphogenesis were significantly associated with excess heritability for COPD (**Fig. 6e**). These findings suggest that risk-associated variants may disrupt normal vascular maturation or vascular response to injury during early life, providing a molecular mechanism linking early-life respiratory insults to increased adult COPD risk and identifying early vascular development as one pathway establishing heritable COPD susceptibility.

## Discussion

This atlas of postnatal human lung development provides an unprecedented analysis of the transcriptional programs that govern lung growth, development, and maturation from birth through adulthood. The dynamic and cell-type-specific transcriptional transitions provide a framework for exploring both fundamental biology and disease mechanisms. We observed both new and previously described cellular interactions, signaling networks, and epigenetic programs that influence the development and maturation of pulmonary structure and function. Furthermore, by mapping out gene expression patterns associated with risk of chronic lung disease, we can now characterize cell type- and temporally specific periods of disease vulnerability, shedding light onto how best to interrogate disease mechanisms (**Fig. 6f**).

By including samples from day 0 of life, within the first hours of the transition to air-breathing, we have established the rapid set of transcriptional programmatic changes that occur as the lung transitions from a high-resistance circulatory system in a placentally supported organism to a low-resistance pulmonary circulation that rapidly assumes the responsibility of gas exchange. The changes observed in the endothelial lineage, and across all cell types, highlight a significant shift in cellular programs that has been difficult to interrogate until now. This provides a new level of understanding of human lung development and the processes that regulate the capacity for respiratory function.

We also identified coordinated downregulation of canonical developmental signaling pathways, including TGF-β, vascular patterning, ECM synthesis, and the IGN, across the postnatal period. Understanding how these pathways and processes are regulated between development and disease could shed insight into how chronic lung disease co-opts developmental pathways during dysplastic repair, and how these processes could be redirected to support lung regeneration.

The lung is not only a respiratory organ but also an immune organ, playing a critical role in host defense, with infections and inflammatory disorders being common occurrences in both acute and chronic lung diseases. Our atlas reveals a striking relationship between the pulmonary endothelium and the immune system. We observed upregulated innate stress and cytokine signaling around birth, followed by a delayed increase in MHC-II gene expression during late childhood. This suggests that endothelial cells may undergo a developmental program that equips them to participate in immune surveillance, potentially linking vascular maturation with the establishment of adaptive immune niches in the lung. Future work should define the functional significance of endothelial MHC-II upregulation and whether it contributes to antigen presentation to T cells, modulates immune cell recruitment, or influences tissue remodeling. Further studies will also need to investigate the role of this axis in chronic lung disease.

We also identified the conservation of age-related developmental signatures between humans and mice. The identification of shared regulatory modules provides a rational starting point for comparative functional genomics to define which pathways are evolutionarily conserved and which are uniquely human. These findings also support the targeted use of murine models to investigate the mechanistic role of shared pathways across species, while simultaneously highlighting the aspects of human lung development that must be modeled in human tissue-specific ways.

Finally, we sought to determine if we could identify time-restricted aspects of vulnerability of adult chronic lung disease. While previous work has identified the cell type of origin for genetic signals, we demonstrate that temporal gene expression during development may also be a critical factor. By incorporating candidate genes identified through GWAS studies across adult lung diseases, we have identified several developmentally regulated expression patterns of genes previously associated with an increased risk of developing chronic lung diseases, such as COPD. These findings support a new approach to understanding disease mechanism and risk. By identifying which cell types and time-periods where risk-associated genes are expressed, we can begin to interrogate mechanisms underlying early life injury risk for adult lung disease. This approach to categorizing disease allows future studies to consider endo-phenotyping disease based on cell type or time period expression of risk variants.

Notably, the heritability signals from our developmental lineage gene sets did not overlap with our mature cell signature analysis, indicating that the RAGs associated with cellular and lineage programs in the mature lung are distinct from the RAGs that are associated with developmental programs. Given the large number of variants linked to chronic lung disease, the mechanistic evaluation of each putative candidate gene to understand its role in conveying disease risk is a lengthy and complex process. This analysis highlights that some of these pathways will need to be studied in the context of lung development. Furthermore, it is well established that chronic lung diseases, such as COPD, are phenotypically complex and likely represent multiple disease entities under one name. By identifying cell-type and time-restricted expression of certain risk-associated genes, this work suggests a cellular, molecular, and genetic framework for endotyping human lung disease.

In summary, we have generated a robust and comprehensive resource that tracks transcriptional programs from birth to lung maturation, revealing several previously unappreciated aspects of human lung development. Given the multiple pathways and cell types that interact to coordinate the complex and delicate growth and maturation of the entire respiratory system, there will be no one-size-fits-all answer for lung regeneration, but this resource for the community provides human-specific data that can be harnessed across many fields. Given the growing need for new therapies for chronic lung disease, the depth and breadth of understanding of lung growth in humans herein establishes pathways and cell types that could be targets of future mechanistic and therapeutic studies. Furthermore, by understanding when and where disease risk is established, we can focus on determining the mechanistic role of associated candidate genes and characterizing putative targeted pathways for both identifying and reducing risk long before the onset of disease.

---

**Supplemental Figure 1S1:**
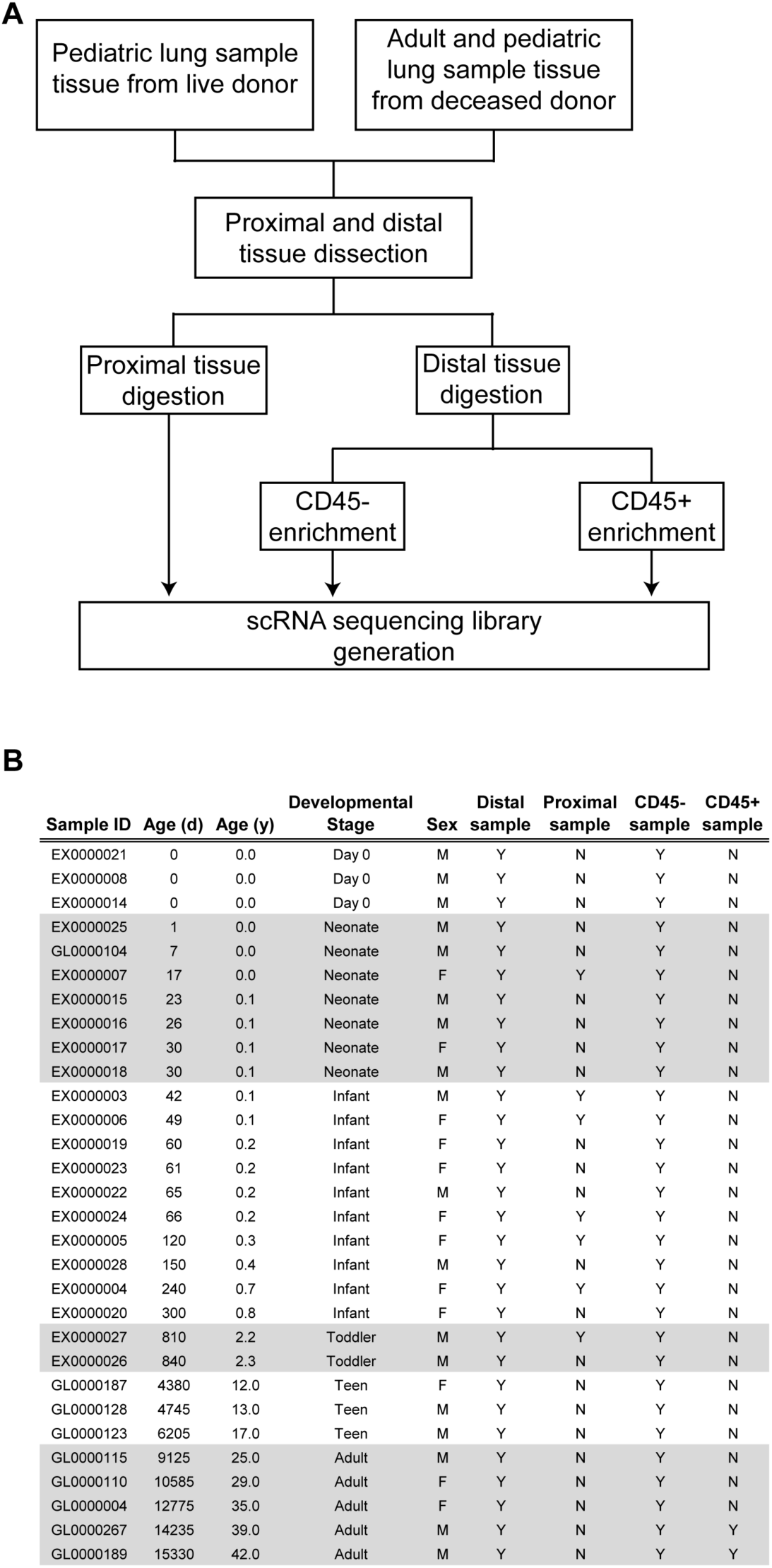
**(a)** Workflow schematic. **(b)** Donor sample characteristics.

**Supplemental Figure 1S2:**
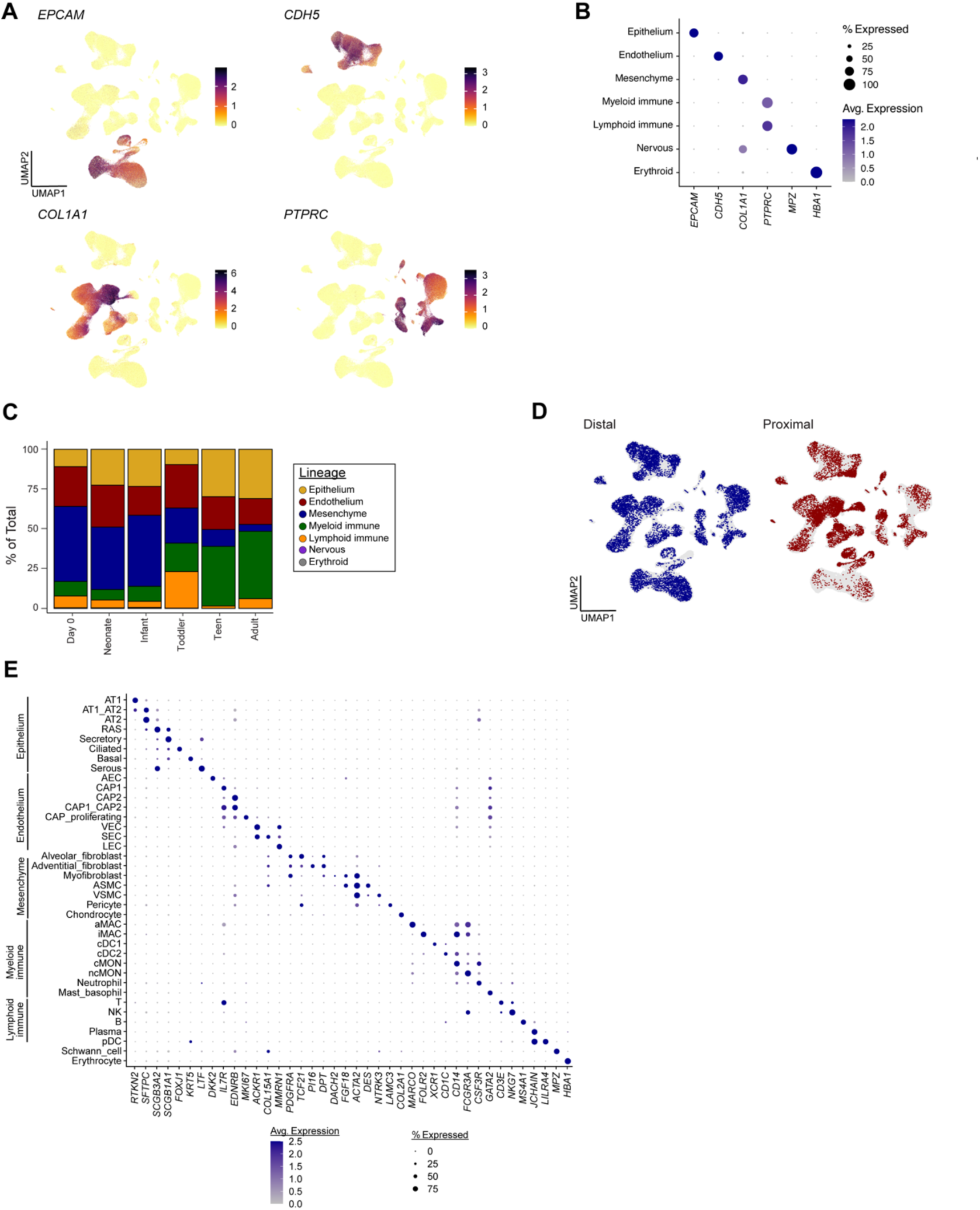
(**a**) UMAP projections of lineage-defining marker genes. **(b**) Dot plot of marker gene expression following lineage assignment. (**c**) Lineage level composition of cells in the dataset. (**d**) UMAP projection of cells obtained from distal and proximal dissections. (**e**) Expression of cell type-specific marker genes.

**Supplemental Figure 1S3:**
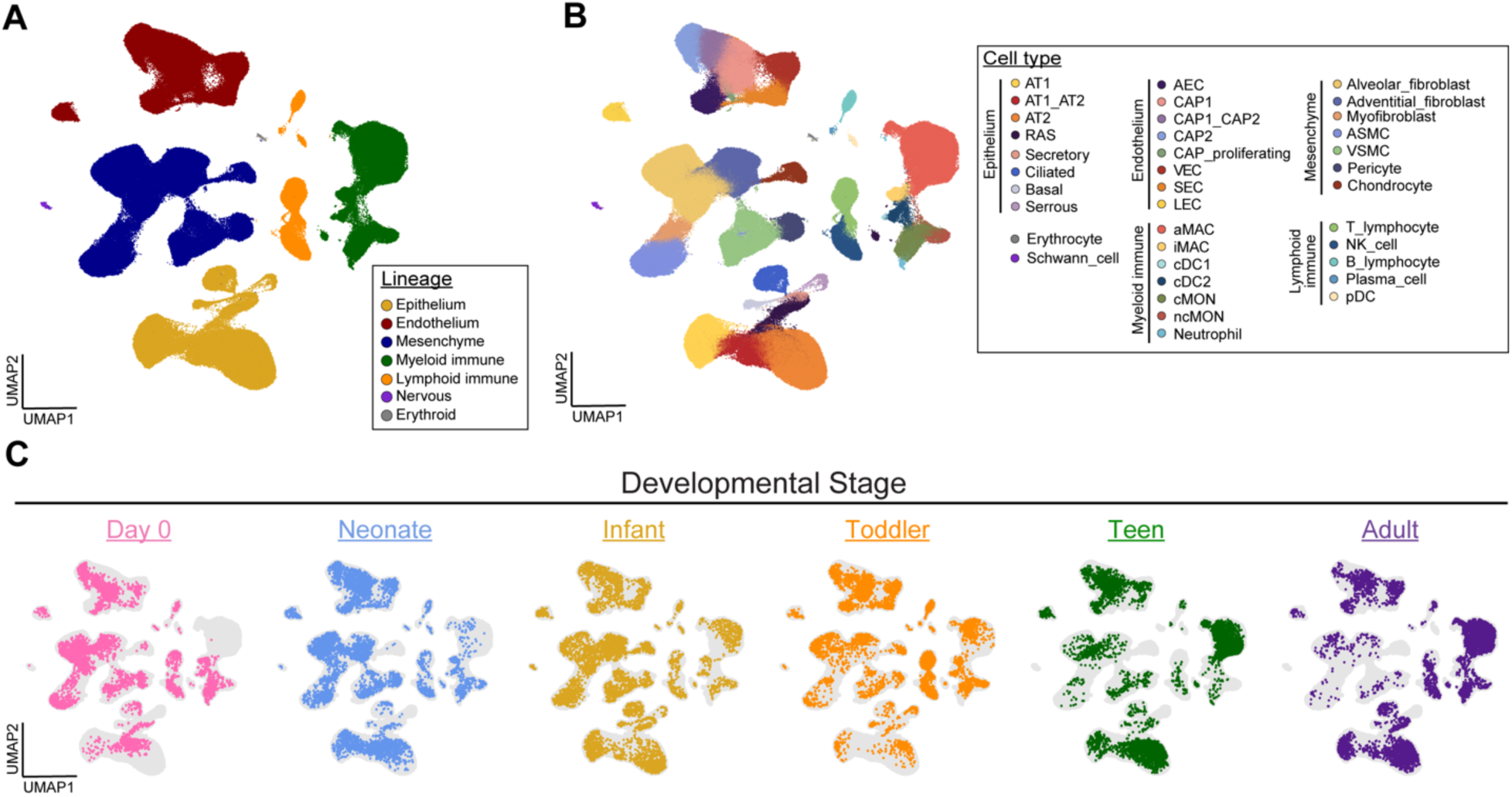
(**a**) UMAP projection of cells from all donors colored by cellular lineage. (**b**) UMAP projection colored by cell type. (**c**) Projection of 5000 cells per developmental stage. Abbreviations: AT1, alveolar epithelial type 1 cell; AT2, alveolar epithelial type 2 cell; RAS, respiratory airway secretory cell; AEC, arterial endothelial cell; CAP1, capillary endothelial 1 cell; CAP2, capillary endothelial 2 cell; VEC, venous endothelial cell; LEC, lymphatic endothelial cell; ASMC, airway smooth muscle cell; VSMC, vascular smooth muscle cell; aMAC, alveolar macrophage; iMAC, interstitial macrophage; cDC1, conventional dendritic cell 1; cDC2, conventional dendritic cell 2; cMON, classical monocyte; ncMON, non-classical monocyte; T, T lymphocyte; B, B lymphocyte; pDC, plasmacytoid dendritic cell.

**Supplemental Figure 1S4:**
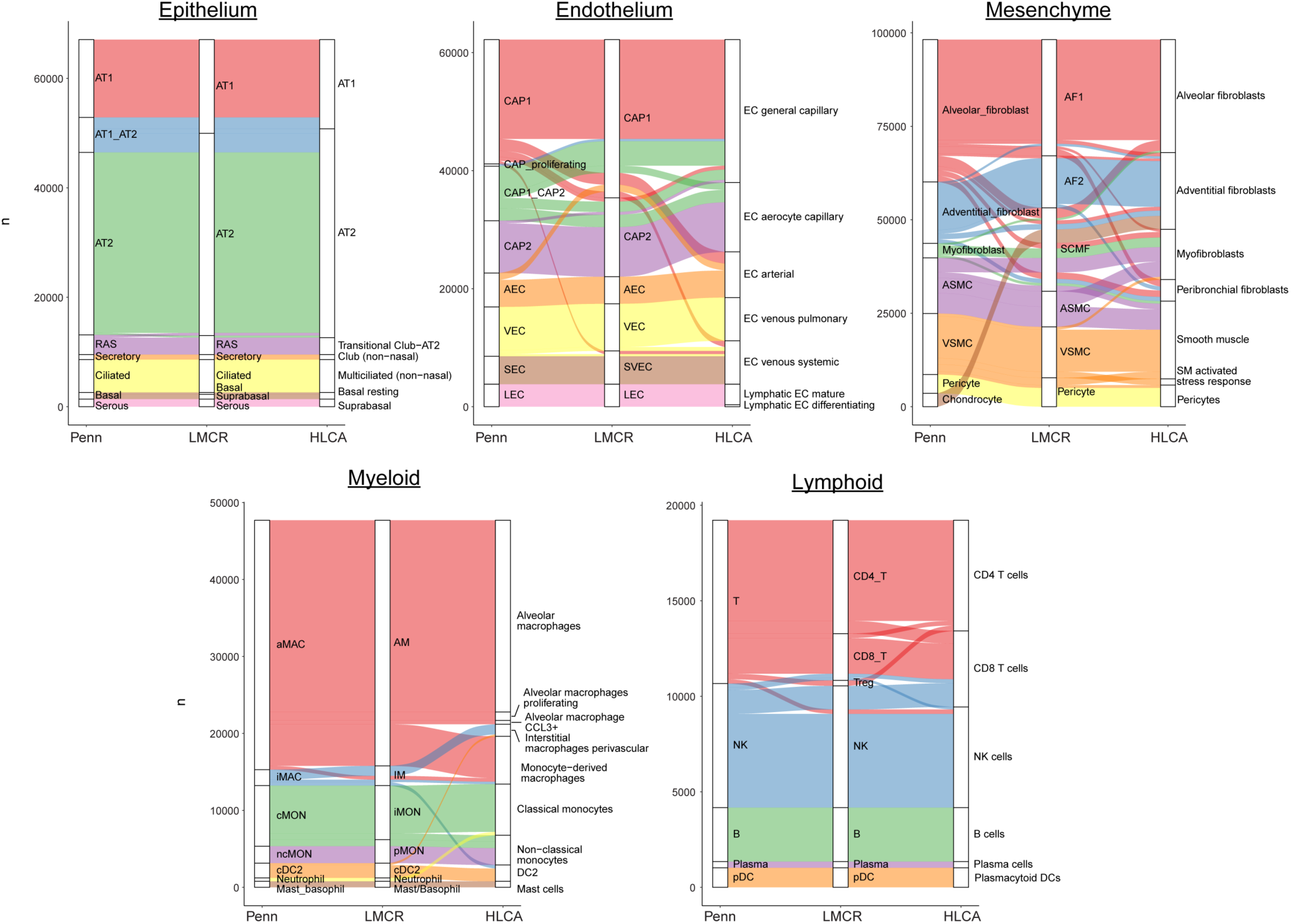
Comparison of cell type annotations between this dataset, LungMAP Cell Reference, and HLCA.

**Supplemental Figure 1S5:**
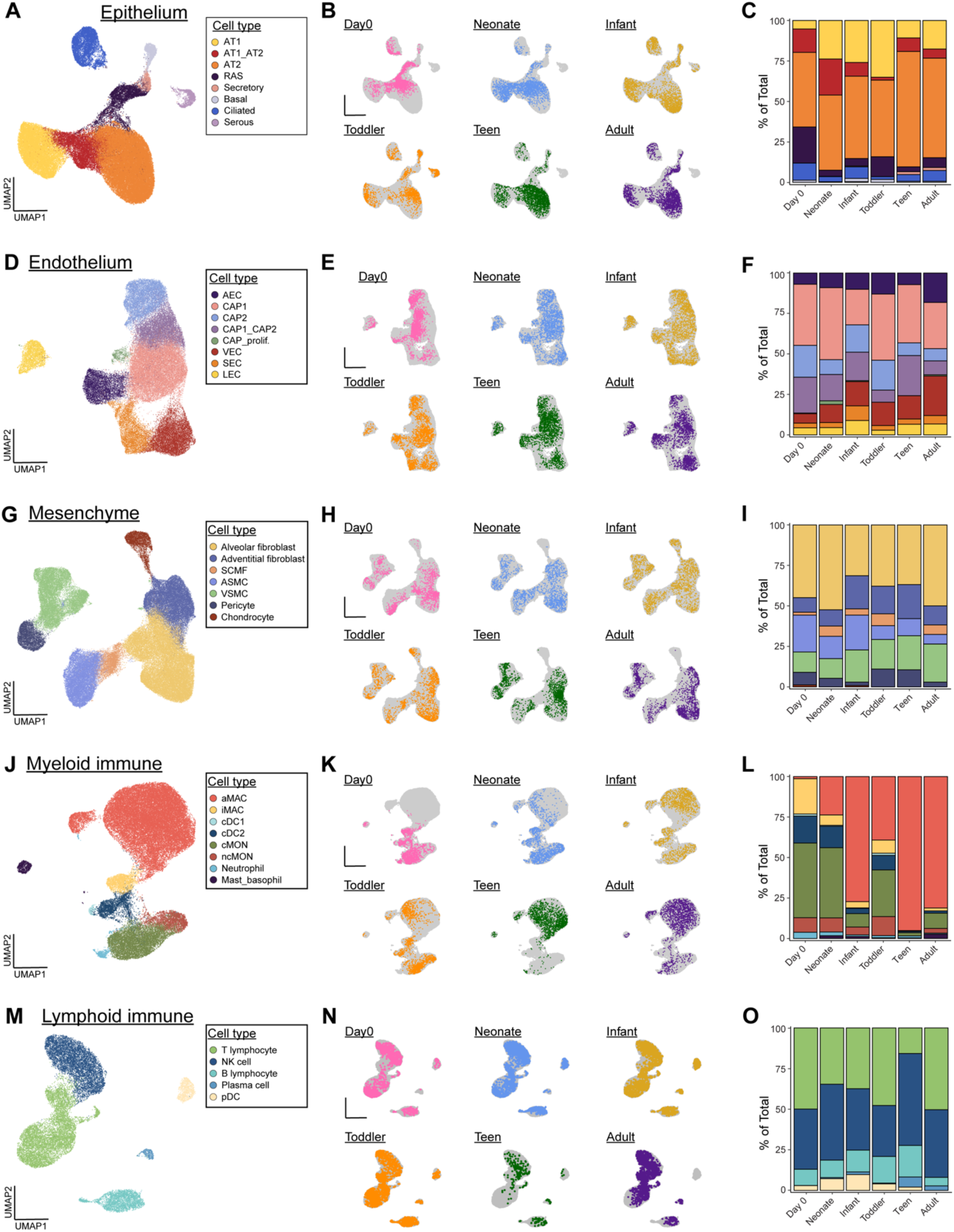
Cell type assignments (**a,d,g,j,m**), UMAP projections for each developmental stage (**b,e,h,k,n**), and composition (**c,f,i,l,o**) for each major cellular lineage.

**Supplemental Figure 1S6:**
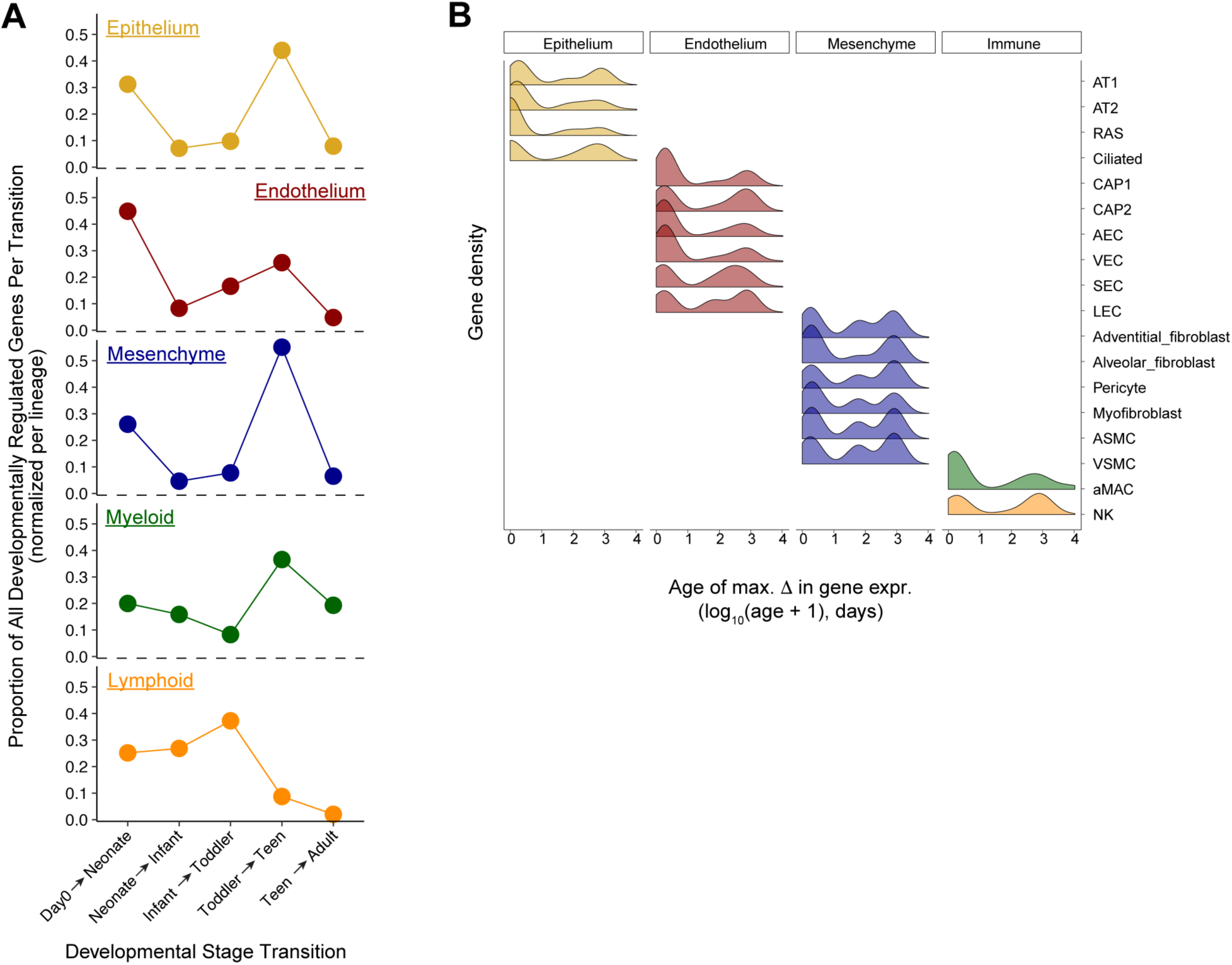
(**a**) DEGs were defined using a categorical approach. The proportion of genes changing between developmental stages are plotted for each lineage. (**b**) Density histogram for major cell types within each lineage.

**Supplemental Figure 1S7:**
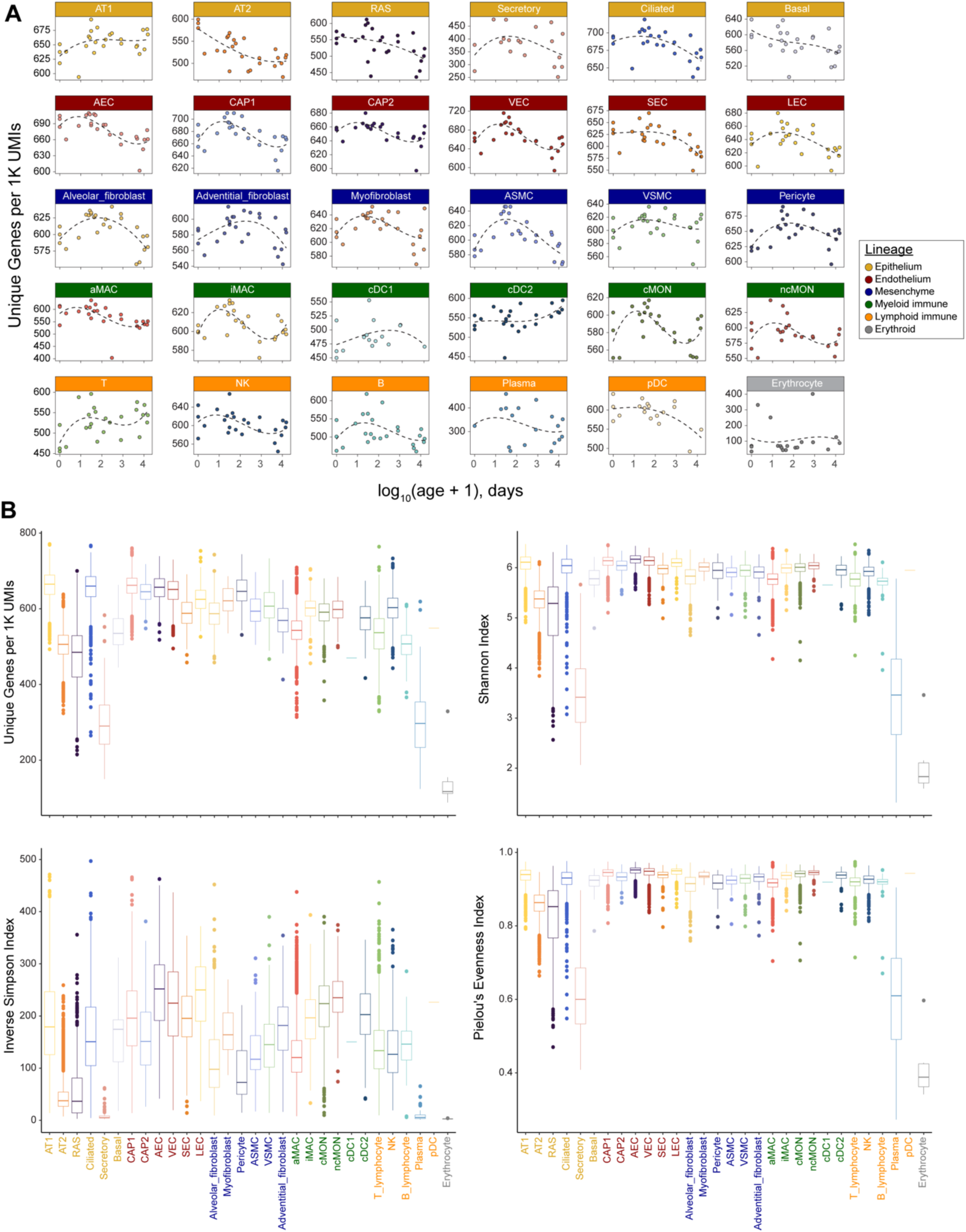
(**a**) Box and whisker plots for each cell type in adult samples displaying unique genes per 1K, Shannon’s index, inverse Simpson’s index, and Pielou’s evenness index. (**b**) The median value of unique genes per 1K per cell type in each sample, plotted over time.

**Supplemental Figure 2S1:**
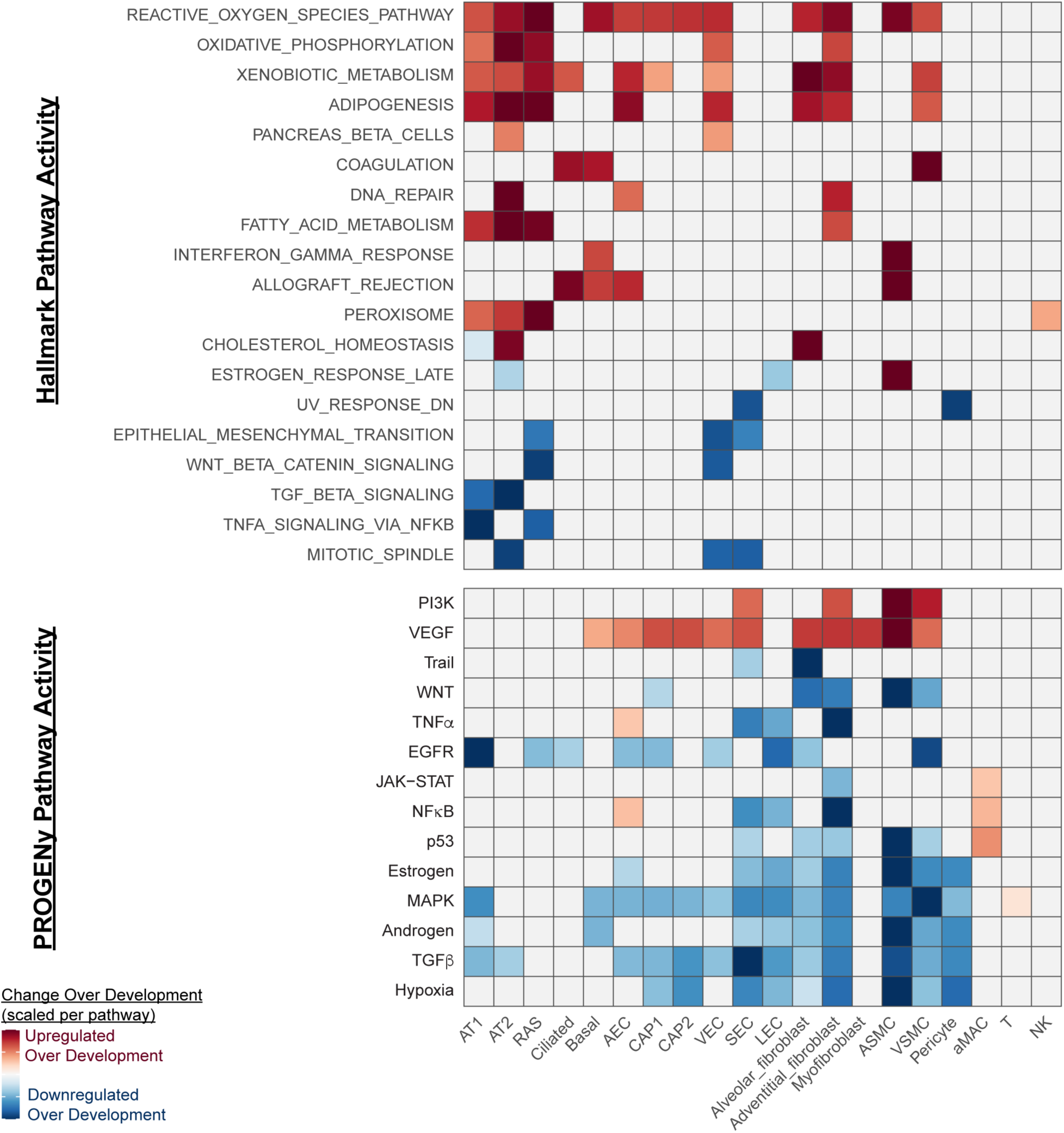
Hallmark and PROGENy pathway score analysis across major cell types. Upregulated pathways are shown in red and downregulated pathways are shown in blue. Scores have been scaled within each pathway.

**Supplemental Figure 2S2:**
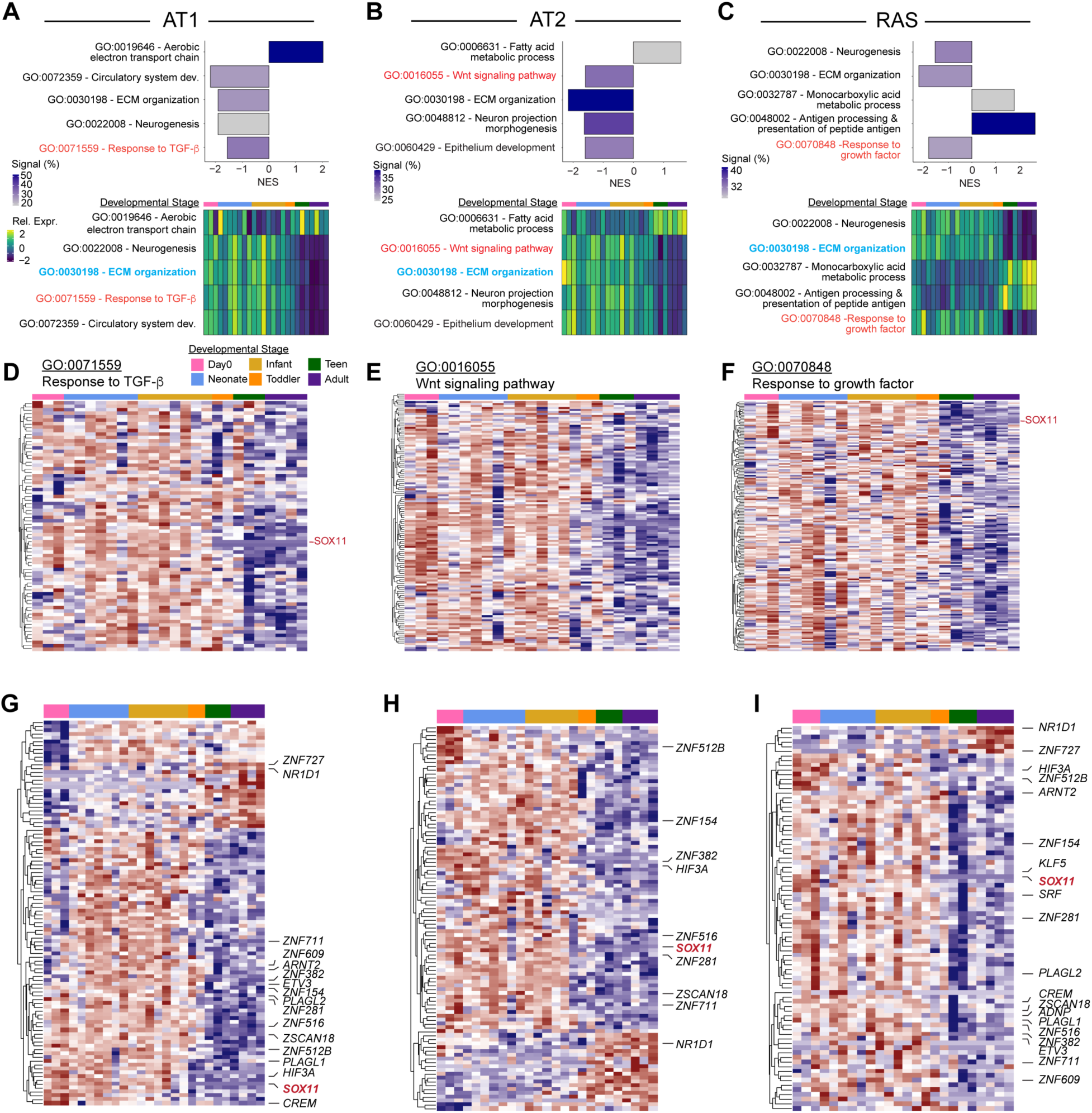
**(a-c)** Gene set enrichment analysis was used to define differentially expressed pathways in AT1, AT2, and RAS cells over time. Net enrichment scores (NES) are plotted for select enriched pathways (Upper). Module scores were generated from the genes in these pathways and plotted from left to right in increasing chronological order (Lower). (**d-f**) Pseudobulk expression values for all genes select pathway are shown. (**g-i**) The transcription factor subset of age-associated DEGs was plotted.

**Supplemental Figure 2S3:**
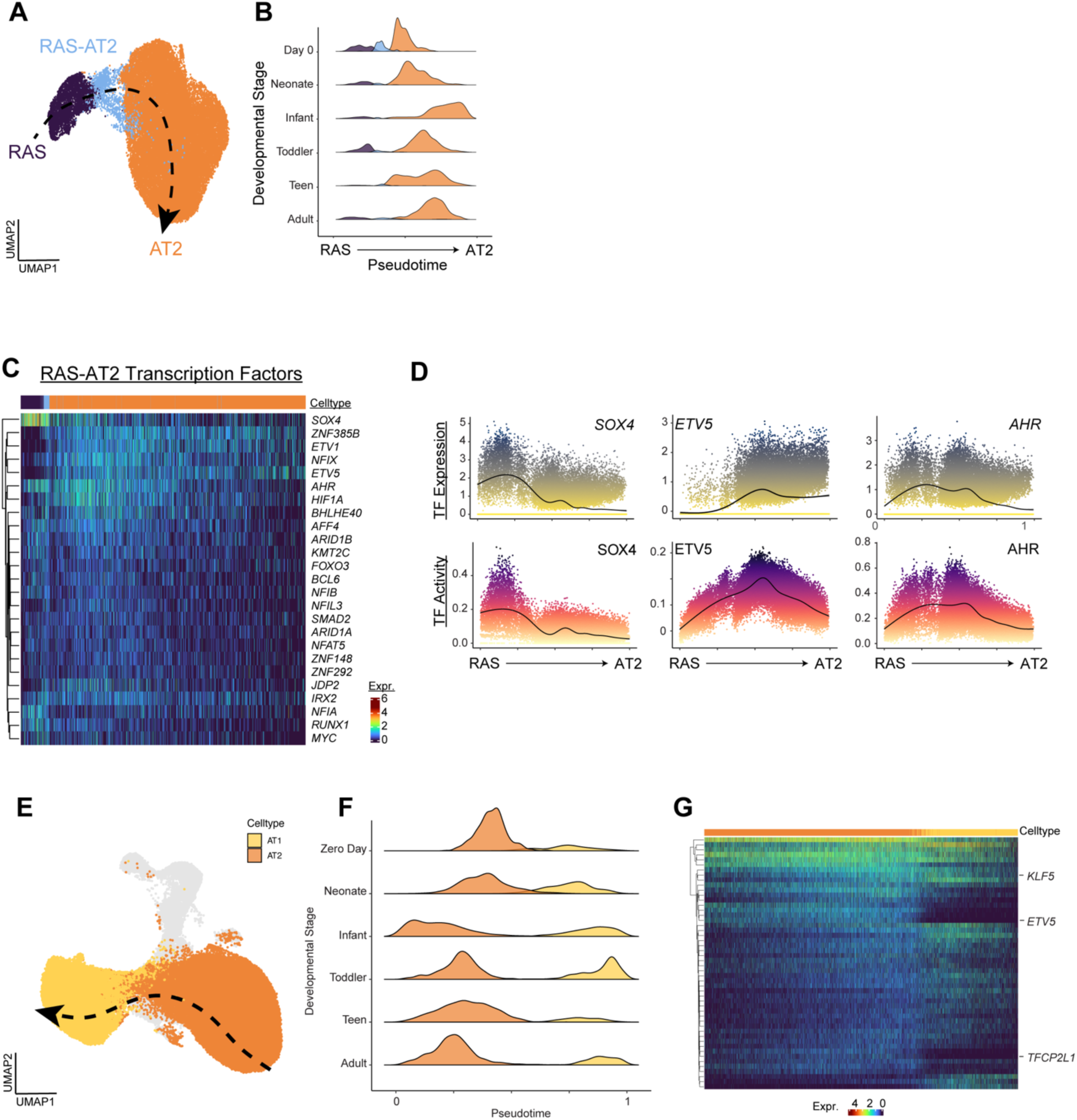
(**a**) RAS-AT2 differentiation was modeled using pseudotime trajectory analysis and (**b**) plotted by developmental stage. (**c,d**) Expression of TFs that varied along the axis of RAS-AT2 differentiation. (**e**) AT2 to AT1 differentiation was modeled as above and plotted by developmental stage (**f**). (**g**) TFs that varied significantly along the axis of differentiation.

**Supplemental Figure 3S1:**
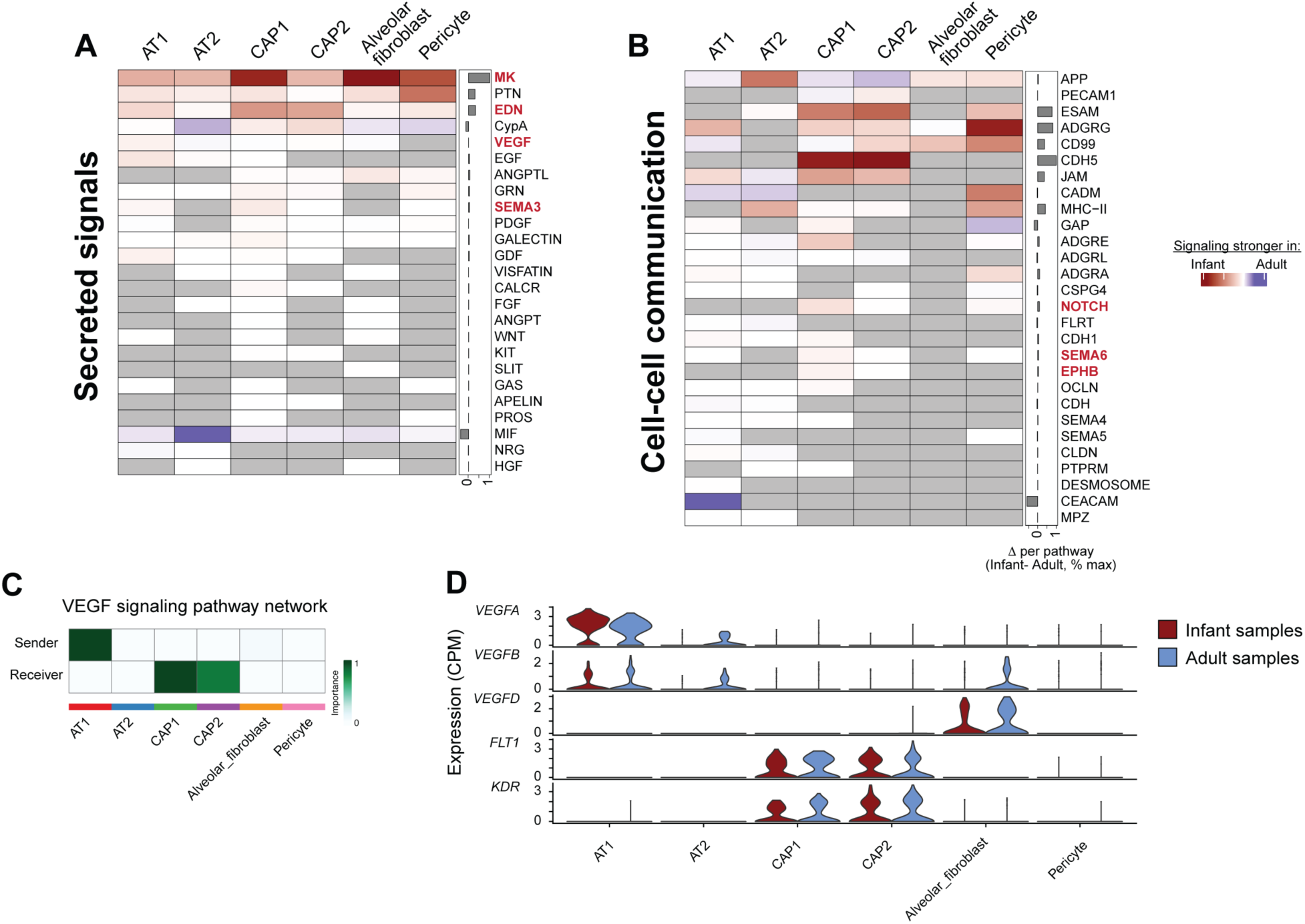
Ligand receptor analysis was used to compare patters of expression for genes in the (**a**) ‘Secreted Signals’ and (**b**) ‘Cell-cell communications’ categories. (**c**) VEGF signaling in cells within the alveolar niche. (**d**) Expression of ligands and receptors in the VEGF signaling pathway.

**Supplemental Figure 3S2:**
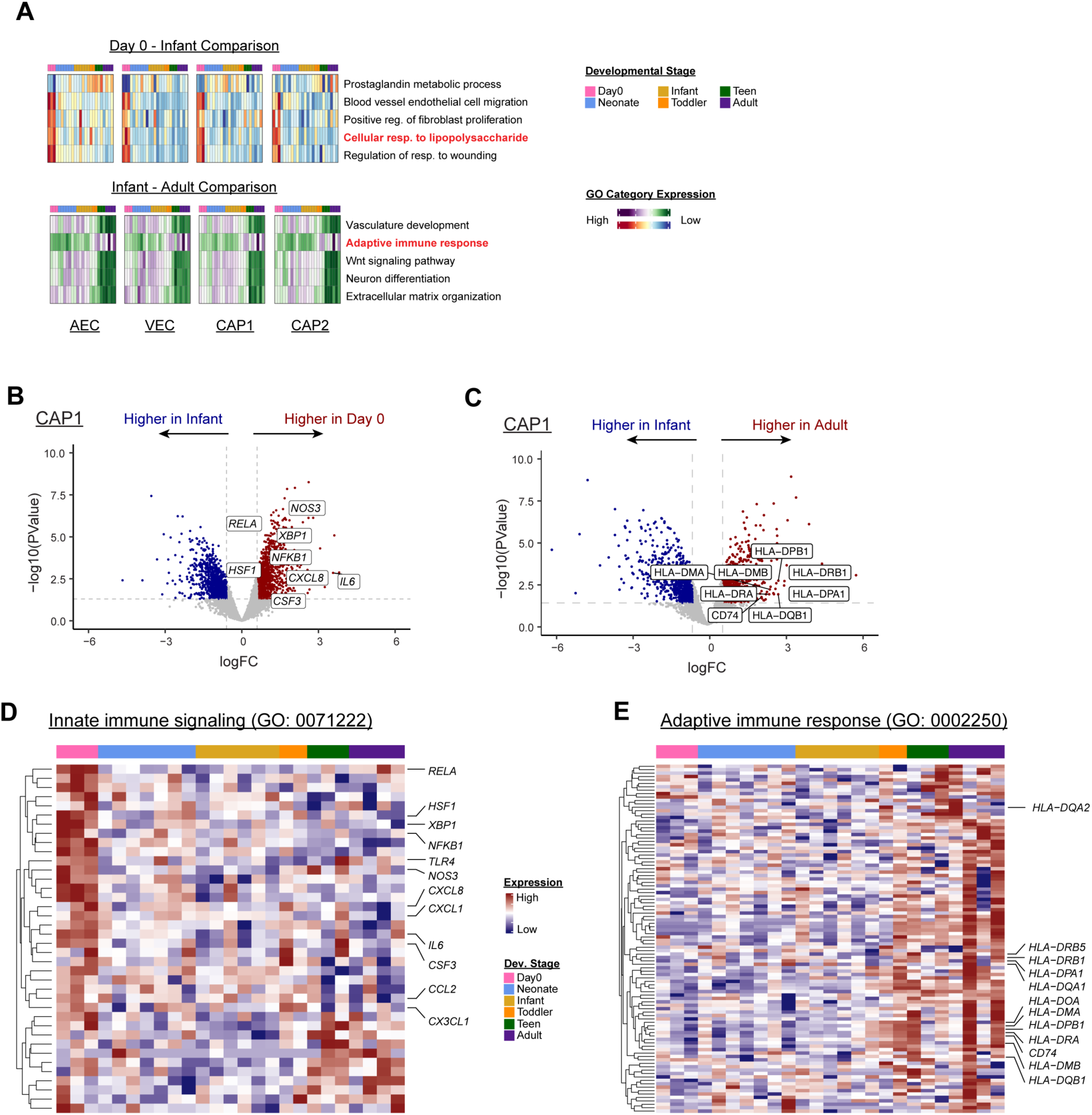
(**a**) GO:BP pathway scores were generated for each of the major pulmonary vascular endothelial cell types and plotted across postnatal age. Volcano plots demonstrating upregulation of immune signaling genes in CAP1 ECs shown in Fig. 3. (**b**) Day 0 neonate vs. infant comparison. (**c**) Adult vs infant comparison. **(d,e**) Heatmaps of genes comprising up- and downregulated GO categories over age.

**Supplemental Figure 3S3:**
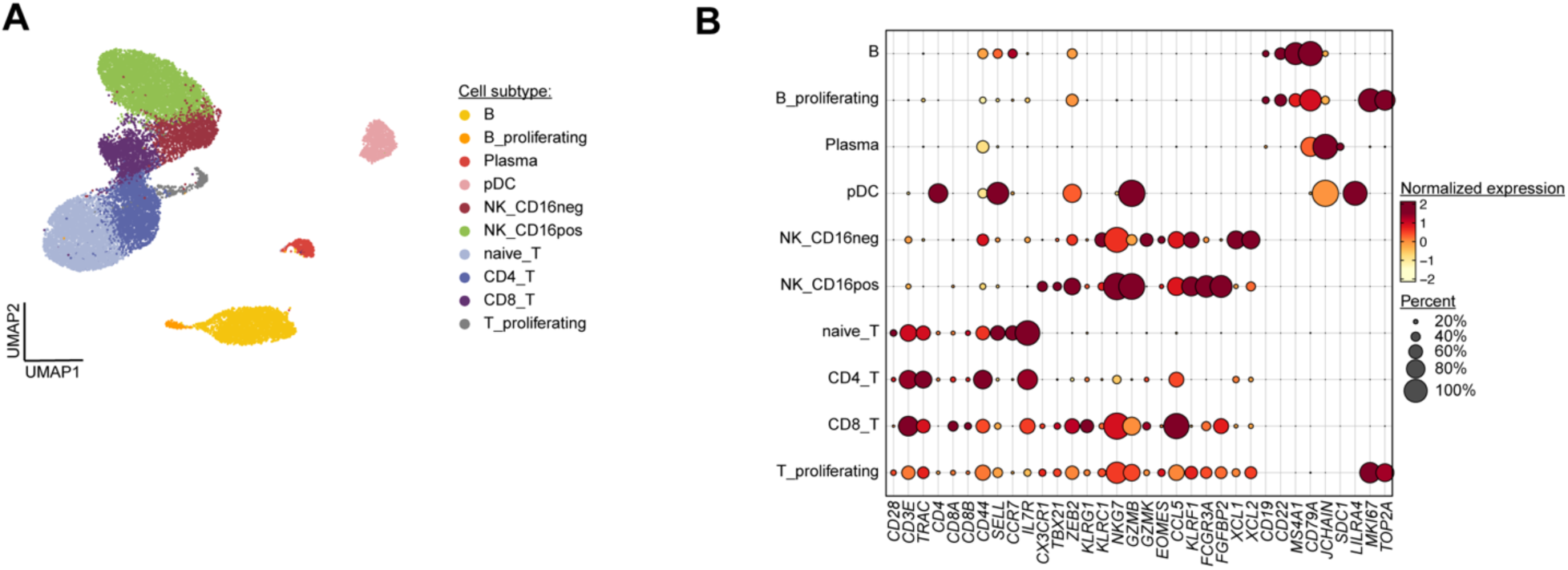
Re-annotation of the lymphoid immune subset after re-clustering. (**a**) UMAP projection of cell populations. (**b**) Marker gene expression.

**Supplemental Figure 4S1:**
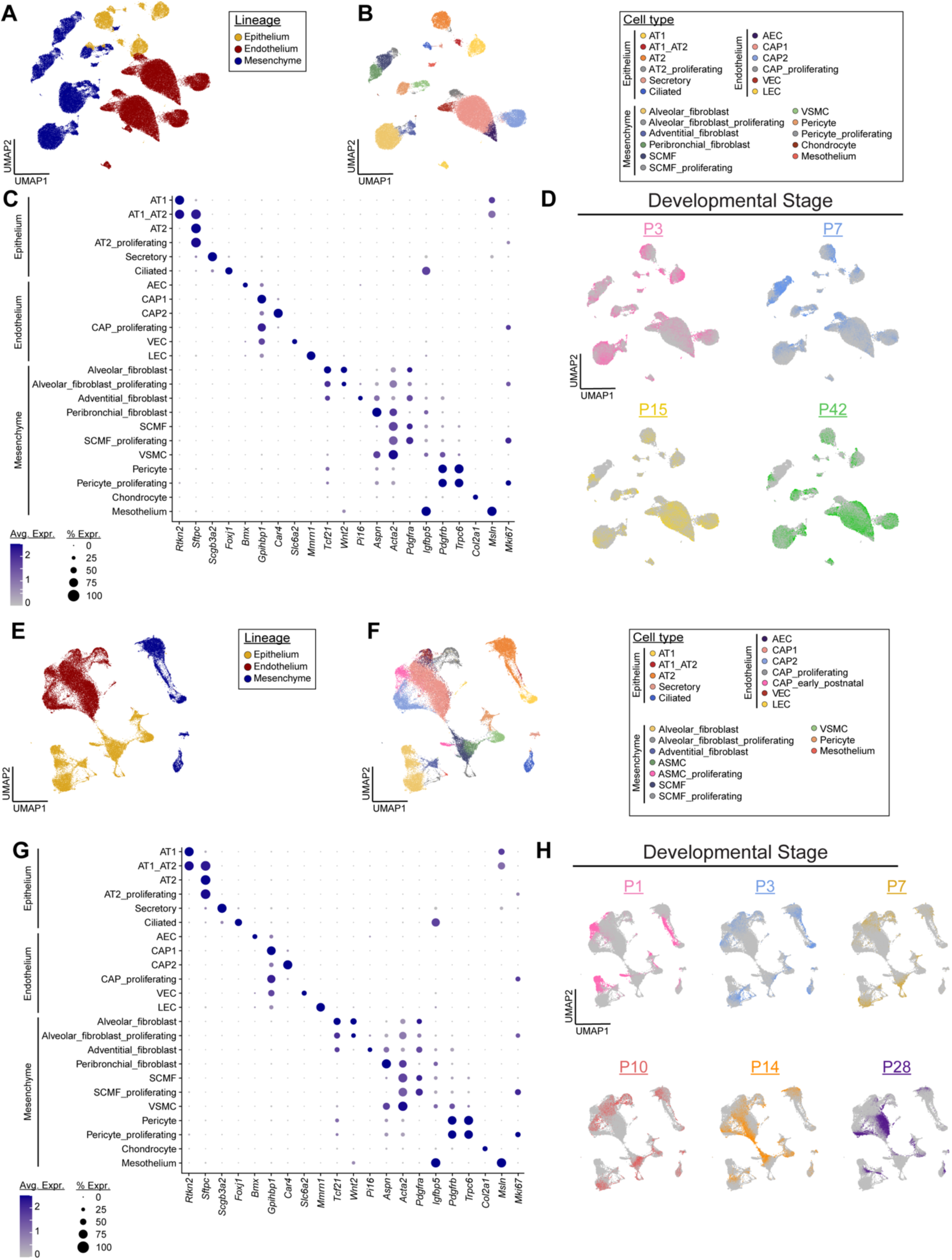
Two existing atlases of mouse postnatal lung development were reannotated using an analogous approach to the human data set. Lineage (**a,e**) and cell type (**b,f**) assignments were performed based on expression of canonical marker genes (**c,g**) and plotted over post-natal age (**d,h**).

**Supplemental Figure 4S2:**
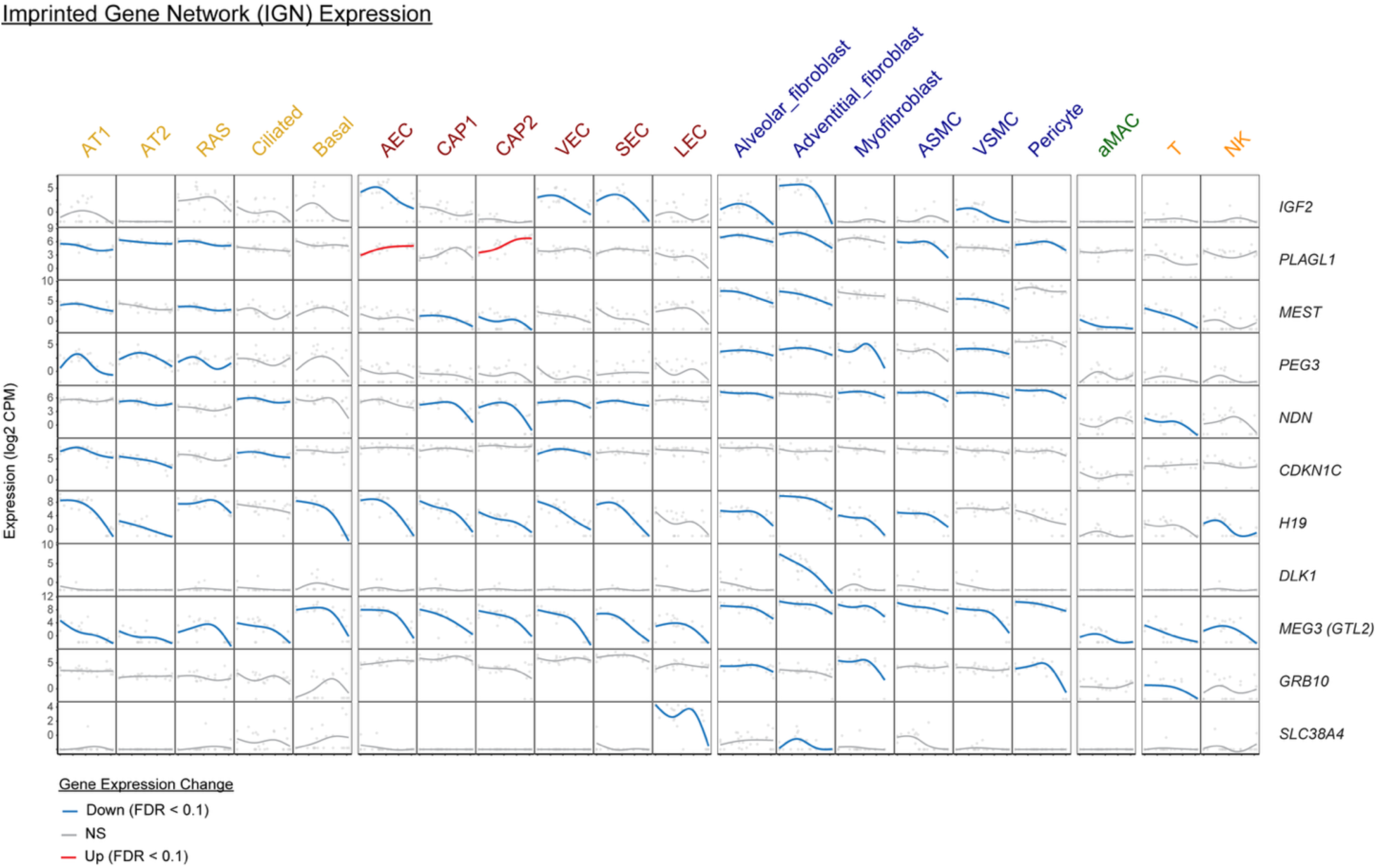
Genes identified as core members of the IGN were plotted over time in major cell types of the distal lung.

**Supplemental Fig. 5S1:**
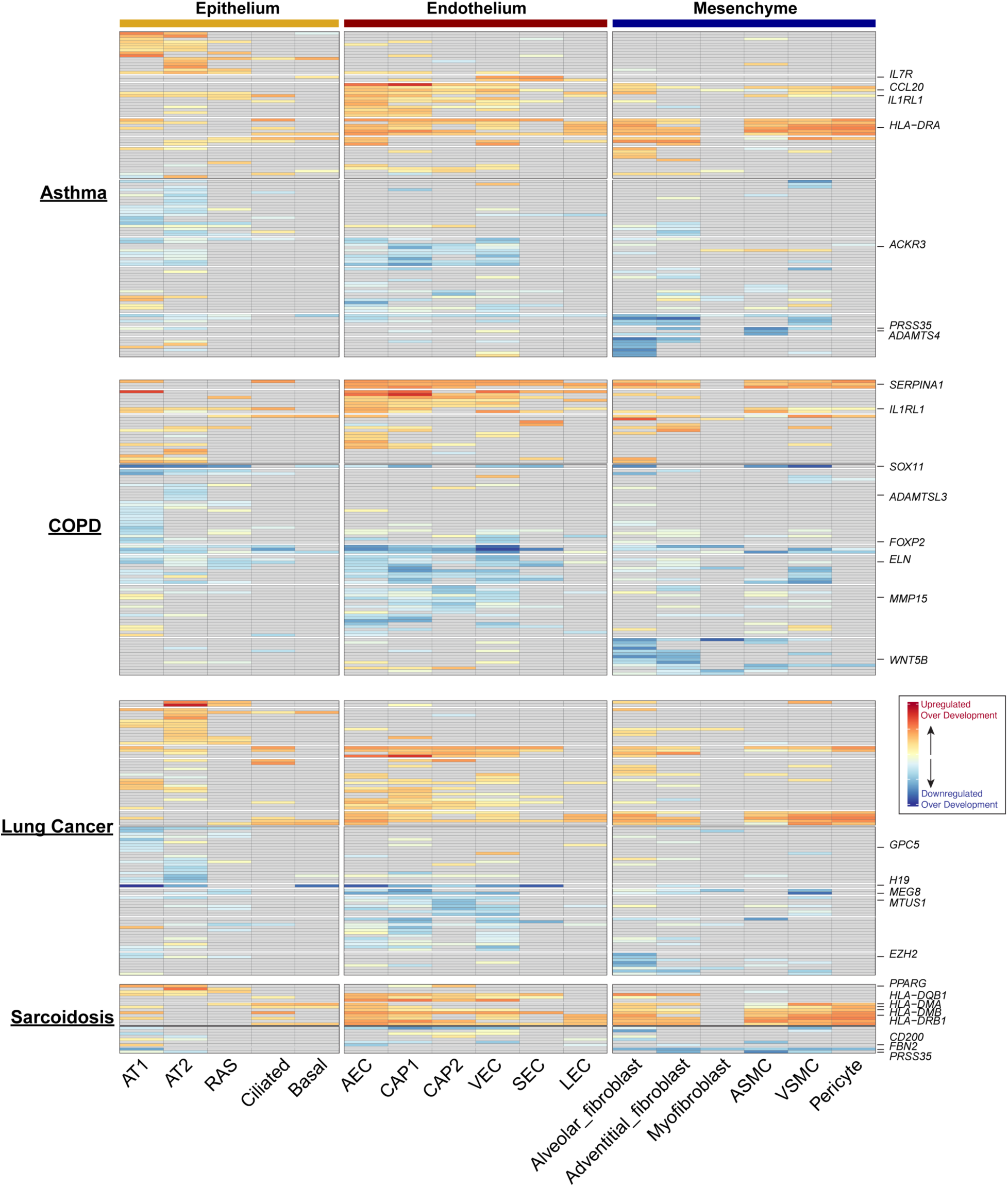
Age-associated RAGs for the major cell types within each disease category.

**Supplemental Fig. 5S2:**
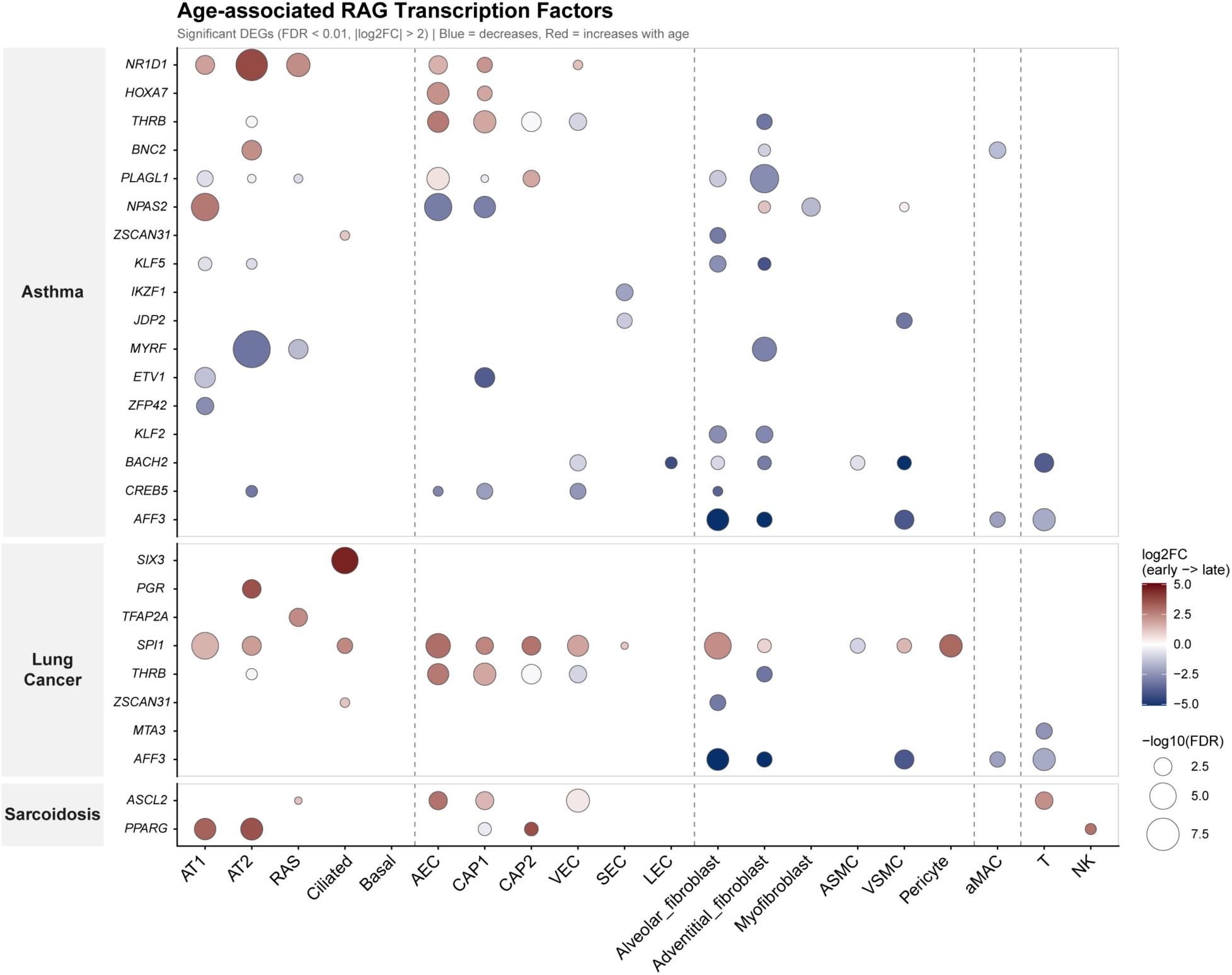
Age-associated RAGs for the major cell types within each disease category (transcription factor subset).

**Supplemental Figure 6S1:**
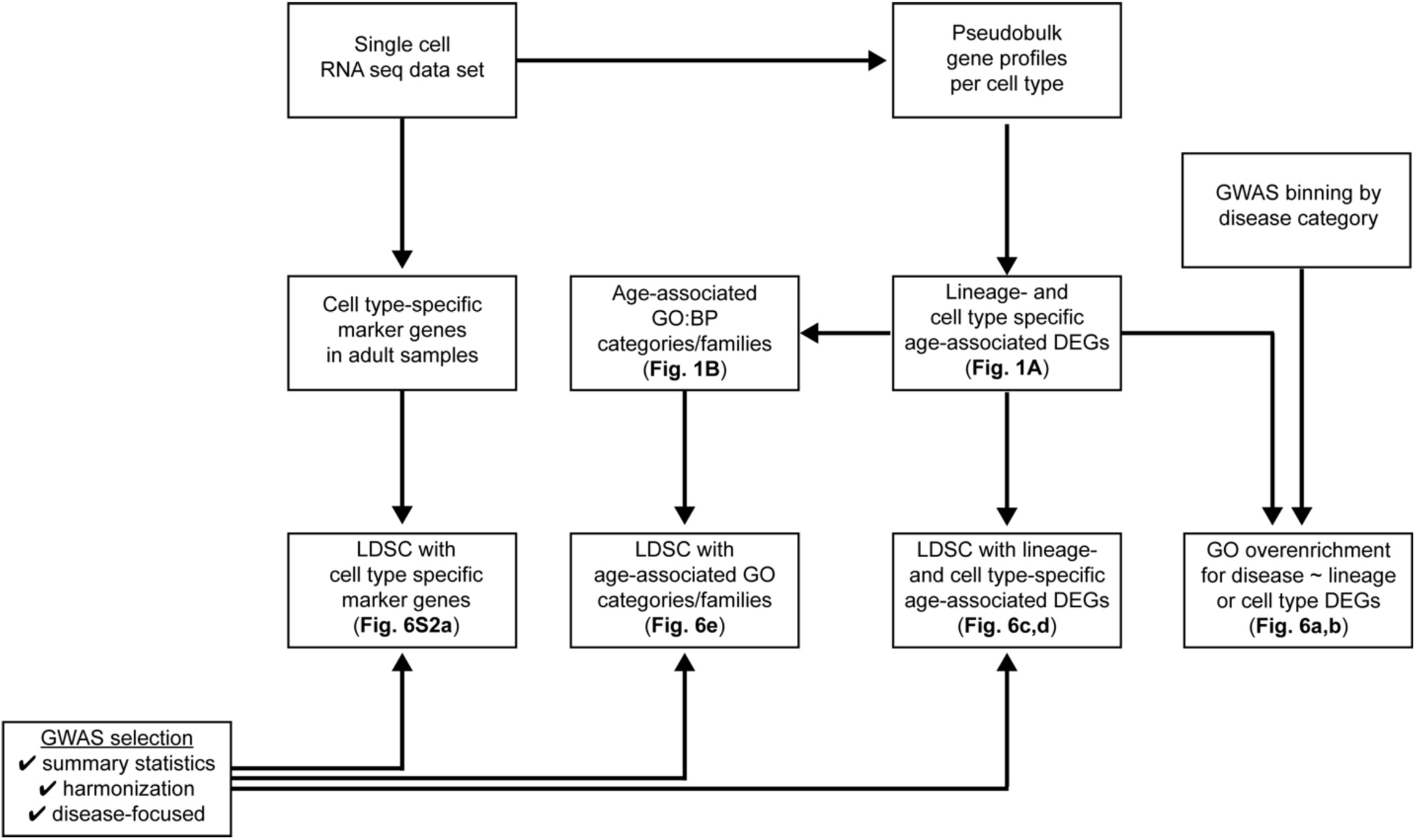
Workflow schematic for GO overenrichment and LDSC analyses

**Supplemental Figure 6S2:**
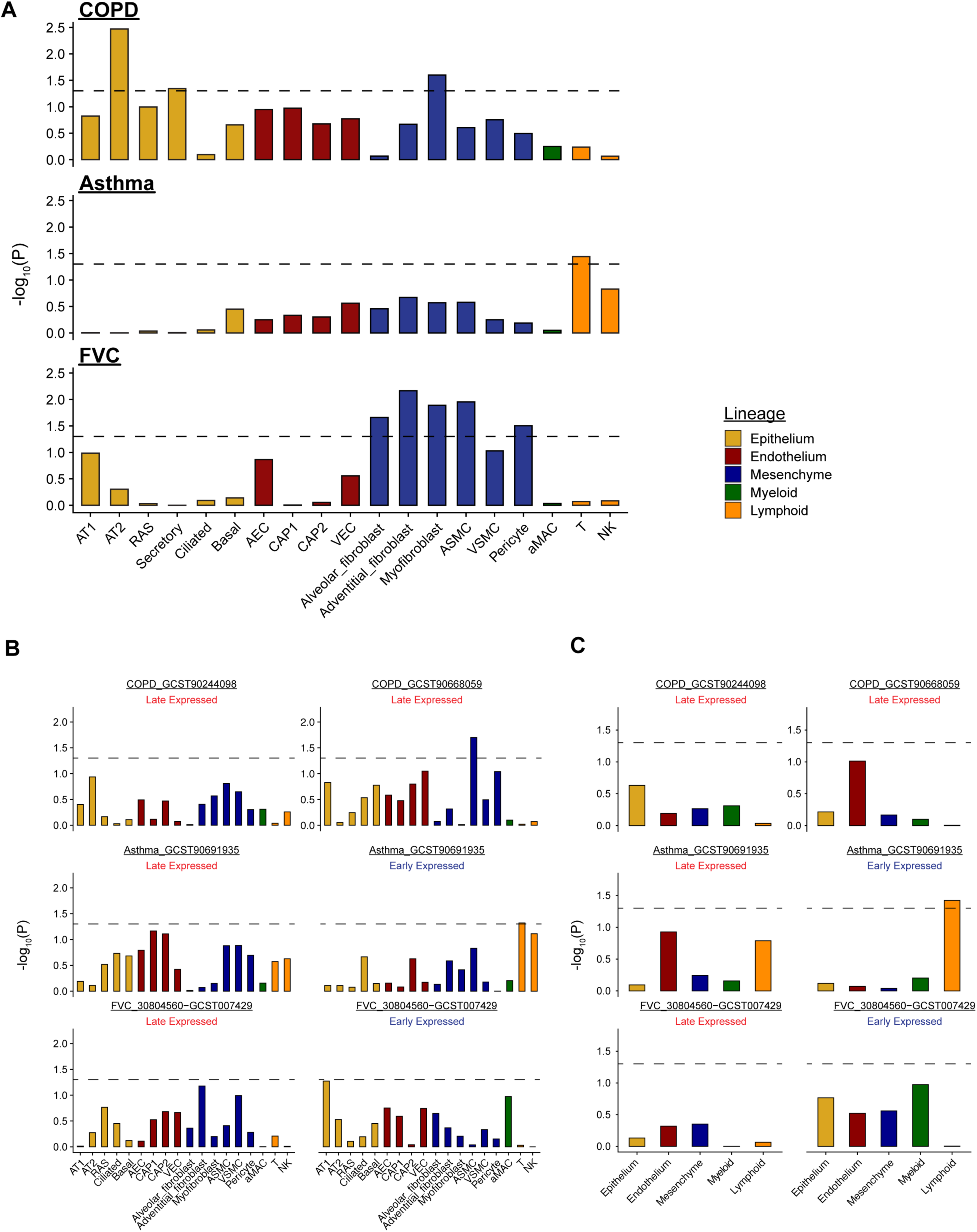
**(a)** LDSC regression was performed at the cell type level using cell type-specific marker genes. Nominal P values are shown with the dashed line denoting P = 0.05. (**b,c**) LDSC regression P values for Early- and Late-expressed gene sets in GWAS for COPD, asthma, and FVC. Age-associated DEG sets were binned at the cell type (**b**) and lineage (**c**) levels.

## Acknowledgments and Funding

The authors would like to acknowledge and thank the core facilities at Penn and CHoP without whom we could not have completed this work. Confocal microscopy was performed in the Penn Cell and Developmental Biology Microscopy Core (RRID: SCR_022373) and the CCR Optical Microscopy and Analysis Laboratory. We also thank the members of the Basil, Morrisey, and Frank labs for their helpful suggestions and discussions over the course of this work.

## Author Contributions

J.D.P: conception and design of the study; acquisition, analysis, and interpretation of data; generation of code and visualization; manuscript preparation

M.P.M.: analysis, and interpretation of data; generation of code and visualization, manuscript preparation

S.C.: acquisition, analysis, and interpretation of data

A.R.P.: acquisition, analysis, and interpretation of data

M.L.: acquisition, analysis, and interpretation of data

Y.Y.: generation of code and visualization

H.H.: preparation of histologic specimens

S.E.S.: acquisition, analysis, and interpretation of data

J.C.: acquisition, analysis, and interpretation of data

S.Z.: preparation of histologic specimens

S.M.L.: sample acquisition, analysis, and interpretation of data P.C.: sample acquisition

L.R.Y.: sample acquisition

E.C.: sample acquisition

W.P.: sample acquisition

D.B.F.: sample acquisition

E.E.M.: conception and design of the study; manuscript preparation

M.C.B.: conception and design of the study; acquisition, analysis, and interpretation of data; manuscript preparation

## Methods

### Human Samples

Lung tissue was obtained either from ‘healthy’ patients at the time of organ procurement or from morphologically normal tissue in pediatric patients undergoing lobar resection of congenital space-occupying pulmonary lesions. Patient samples were collected under the protocol “Molecular and Clinical Analysis of Syndromic and Isolated Anomalies of the Foregut” that has been approved by the Institutional Review Board of the Children’s Hospital of Philadelphia (IRB protocol, 12-009788). Adult samples were collected as previously described^23^.

### Tissue preparation and magnetic-activated cell sorting

De-identified human lung tissue was obtained and processed as previously described^65^. Briefly, samples consisting of up to 8g of human lung tissue were dissected and manually disrupted with scissors. Tissue was resuspended in digestion buffer, transferred to GentleMACS C tubes, and incubated at 37°C for 35 minutes with intermittent agitation. Digested tissue was filtered through a 70 μm Falcon cell strainer, resuspended in ACK lysis buffer and then incubated for 5 minutes at room temperature. Cells washed once with FACS buffer and filtered through a 40 μm cell strainer. Viability was assessed using Countess II (Lifetech) in conjunction with trypan blue exclusion. Cells were washed once and resuspended in 0.04% bovine serum albumin (Jackson) in Dulbecco’s phosphate buffered saline for downstream processing.

### Single-cell RNA Sequencing Library Preparation

*Single*-cell transcriptome libraries were prepared using the Chromium Single Cell 3′ Gene Expression v2/v3 platform (10x Genomics) following the manufacturer’s instructions. Reverse transcription, cDNA amplification, and library construction occurred according to standard protocols. Libraries were indexed, pooled, and sequenced on Illumina NovaSeq platforms to a target depth of 20,000–30,000 read pairs per cell.

### Read Alignment, UMI Processing, and Gene Quantification

Raw sequencing data (BCL files) were converted to FASTQ format using bcl2fastq. Reads were aligned to the GRCh38-2020A human reference genome using STARsolo (STAR v2.7.9a), which performed alignment, barcode and UMI correction, and gene-level quantification in single-cell mode^106^. STARsolo was run with 10x Genomics Cellranger chemistry presets and configured to output both exonic gene counts (--soloFeatures Gene). Only barcodes classified as valid cells by STARsolo’s Emptydrops implementation were included. Cells with fewer than 1200 UMIs, cells exceeding three median absolute deviations (MADs) above the gene-count distribution, and cells with more than 20% mitochondrial RNA content were excluded. Potential doublets were identified using Scrublet and excluded. All remaining cells were retained for downstream integration and clustering.

#### Data Integration Using scVI

Count matrices were converted into AnnData format using sceasy, and highly variable genes were identified using Seurat’s variance-stabilizing transformation. Genes associated with mitochondrial, ribosomal, and MHC loci were excluded from the HVG set to prevent technical factors from driving the latent space. Integration was performed using scVI (scvi-tools 1.2.1), which models gene counts using a deep generative framework^107^. Models were trained with donor identity as the batch variable and a latent dimensionality set to 30 and number of layers set to 2 and library platform was used as a covariate. After training, the scVI latent representation was used for clustering, visualization (UMAP), and downstream analysis.

#### Cell Type Annotation

Clusters were annotated as previously described using established canonical markers for major lung lineages, including epithelial, endothelial, mesenchymal and immune populations^108^. Initial marker-based annotations were reviewed manually to ensure biological consistency and to resolve ambiguous clusters. To further refine cell identities and ensure concordance with existing lung atlases, label transfer was performed using Seurat’s reference-mapping framework with both the Human Lung Cell Atlas (HLCA)^13^ and the LungMAP CellCards (LMCR) reference^14^. Transferred labels were integrated with marker-based assessments to generate the final, high-confidence cell type annotations. Alluvial plots comparing nomenclature consistency were generated using the ‘ggalluvial’ package.

### Pseudobulk Aggregation and Differential Gene Expression Analysis

To perform robust differential gene expression (DGE) analysis across developmental time, a pseudobulk framework was used with single-cell counts aggregated at the donor-cell type level. For each annotated cell type, all cells originating from the same donor were summed to create a donor-level expression profile, using the Seurat2pb function implemented within the edgeR package^109^. Only donor-cell type combinations containing high-quality cells were retained to ensure adequate biological representation and statistical stability. Pseudobulk analyses were then carried out using edgeR, which provides a well-established model for overdispersed count data and appropriately treats donors as the units of replication.

### Developmental Time Modeling

Developmental age for each donor was transformed using log10 scaling to reflect the nonlinear progression of early postnatal lung maturation. Gene expression was modeled as a smooth function of this log10-adjusted age using natural spline regression with three degrees of freedom.

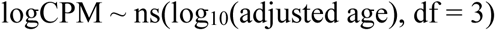

For each gene, edgeR’s generalized linear model framework was used to fit quasi-likelihood (QL) models in which logCPM expression was regressed against the spline basis while including a donor-level intercept term. This spline-based approach accommodates both gradual and rapidly changing developmental expression patterns without imposing linearity. Samples generated using the 10X version 2 chemistry were removed from pseudobulk analyses due to excessive technical variance. Significance of developmental trends was assessed using edgeR’s quasi-likelihood F-test, comparing the full spline model with a reduced intercept-only model. P-values were corrected for multiple testing using the Benjamini-Hochberg method, and genes with a false discovery rate (FDR) below 0.05 were considered significantly associated with developmental age. Genes passing this threshold were subsequently classified according to their fitted spline shapes to identify increasing, decreasing, transient, or more complex expression trajectories across pediatric lung development.

### Developmental Gene Clustering

All genes identified as significantly associated with developmental age were grouped into transcriptional trajectories using k-means clustering (k = 6), allowing us to define major developmental expression patterns across the pediatric lung. These cluster-level temporal profiles were visualized using the ComplexHeatmap R package^110^, which provides a flexible framework for generating high-resolution, annotated heatmaps suited for large gene sets. To further interpret the biological significance of each gene cluster, we performed functional enrichment analysis using the clusterProfiler package^111^, enabling systematic evaluation of Gene Ontology (GO), KEGG, and other pathway-level enrichments associated with each developmental trajectory.

### Alpha diversity measurements

Alpha diversity indices were calculated using the ‘vegan’ package in R after evenly downsampling to 1000 reads per cell. Plotted values represent the median count per donor-cell type combination.

### Regulon/Transcriptional Activity

Transcription factor activity was inferred using the pySCENIC workflow^112^. Co-expression modules were first identified with GRNBoost2 and refined into high-confidence regulons through motif enrichment analysis using human cisTarget databases. Regulon activity was then quantified with AUCell, which scores the enrichment of each regulon within a cell’s ranked expression profile. The resulting regulon activity matrix was used to compare TF-driven regulatory programs across developmental stages. Transcriptional activity was also assessed using the AddModuleScore function in Seurat or Gene Set Variation Analysis^113^ in conjunction with gene sets derived from the Molecular Signatures Database^114,115^. Pathway analysis was also performed on pseudobulk expression profiles using the ‘progeny’ package^116^ in R.

### Trajectory Analysis

Trajectory analysis was performed using the ‘slingshot’ package in R^117^ as previously described^108^. Briefly, cells were re-clustered and a supervised approach was used to generate a trajectory originating in RAS cells and terminating in AT2 to evaluate the putative RAS-AT2 trajectory. Gene expression along this trajectory was evaluated using the ‘tradeSeq’ package^118^.

### Ligand Receptor Analysis

Ligand receptor analysis was performed using the ‘CellChat’ package in R^119^ as previously described^108^. Cell types with adequate cellular representation were evenly downsampled to 100 cells per developmental stage. For analyses presented in the paper samples in the ‘Infant’ age bin were compared to samples in either ‘Day 0’ or ‘Adult’ age bins. Pathway signaling strength matrices were extracted from the cellchat object and used for downstream plotting.

### Evaluation of conserved cross-species patterns of age-associated transcriptional regulation

Data from Zepp et al.^58^ and Guo et al.^57^ were re-clustered using only postnatal samples. Cell annotation was performed as described previously^108^. Beta coefficients were extracted from gene-specific linear models to assign directionality, and the subset of genes that were concordantly regulated and statistically significant in both mouse data sets was identified. Human orthologs were identified for each gene, and the set intersection of human and mouse developmentally regulated genes was identified for each cell type.

### Chronic lung disease GWAS parsing

The NHGRI-EBI Catalog of human GWAS was downloaded and parsed locally. Broad disease category bins were created and refined using regular expression parsing. Age-associated DEGs were intersected with genes present in the ‘MAPPED_TRAIT’ bin and used for downstream associations. Patterns of gene expression were defined using the following logic: if significantly age-associated in 1 cell type, annotated as “celltype_specific”; if significantly age-associated in 1 lineage and >1 cell type, annotated as “lineage_specific”; if significantly age-associated in >4 cell types and > 1lineage, annotated as “pleiotropic.”

### Partitioned heritability analysis

To evaluate whether genetic risk for disease is enriched near genes with cell-type–specific expression changes, we performed stratified LD score regression (S-LDSC). Differential expression analyses were conducted within each cell type using pseudobulk expression profiles generated for each donor. Gene expression counts were aggregated by cell type and donor, and differential expression between early and late samples was evaluated using a generalized linear modeling framework. For each cell type, genes significantly upregulated or downregulated with age were identified and used to define cell-type–specific gene sets.

For each gene set, genomic annotations were generated by assigning SNPs to genes with a ±50 kb window around the transcription start and end sites. Binary annotation files were created for each chromosome using the LD Score Regression software package. LD scores were then calculated using European reference genotypes from the 1000 Genomes Project (Phase 3) with the recommended 1 cM window.

GWAS summary stats were downloaded from the NHGRI-EBI catalog for the following traits COPD (GCST90668059), FVC (GCST007429) and Asthma (GCST90691935). GWAS summary statistics were first processed using the LDSC munge_sumstats.py utility to harmonize alleles and restrict analyses to HapMap3 SNPs.

Partitioned heritability analyses were performed using stratified LD score regression implemented in LDSC (v1.0.1). For each cell-type–specific gene set, SNP heritability enrichment was estimated using the cell-type–specific analysis framework (--h2-cts). The baselineLD v2.2 annotation model was included as a control to account for known functional genomic features, including coding regions, conserved elements, regulatory annotations, and chromatin marks. To control for general enrichment near expressed genes, we additionally included a background annotation consisting of all genes detected in the single-cell RNA-seq dataset. Statistical significance of annotation coefficients was assessed using block jackknife standard errors.

### Immunofluorescence (IF) Imaging

Tissue was obtained from lungs as above and preserved in 2% or 4% paraformaldehyde (PFA) overnight. Tissue samples were then washed with PBS and then dehydrated to 100% ethanol over the next 24 hours, before paraffin embedding. Histologic sections were prepared from tissue blocks and deparaffinized for IF or RNAscope imaging. The following antibodies were used for IF staining: human HLA-II (rabbit, Thermo Fisher cat. RBM1-2967-P0, 1:100); VE-Cadherin (mouse, Thermo Fisher cat. 5012896, 1:100), LAMP3(rat, Novus cat. DDX0191P-100, 1:100), IGF2BP3/IMP-3 (mouse, Thermo Fisher cat. 66247-1-Ig,1:200).

### RNAscope Analysis

For RNAscope, samples were preserved and prepared as described above. RNAscope was performed using the Fluorescent Multiplex Reagent kit v2 (ACDBio). Probes were purchased from ACDBio and included: GPIHPB1 (cat. 1811681-C3, C3), H19 (cat. 400771, C1), PDGFRA (cat. 604481-C3, C3), MEST (cat. 467121, C2), SOX11 (cat. 443671, C2), AGER (cat. 4710121, C1), LAMP3 (cat. 468761, C1), SCGB3A2 (cat.549951, C1).

### Confocal microscopy and image processing

RNAscope and IF images were captured using a Leica Stellaris 5 confocal microscope. Z stacks of 40- or 63X zoom were taken and image processing was performed using the FIJI software suite.

### Data and code availability

Single-cell RNA-sequencing data have been deposited at GEO as GSE324191and are publicly available as of the date of publication. The scRNA-seq datasets can be viewed online in an interactive browser (bit.ly/3PzHp04). All code used to generate the data in this manuscript is adapted from base R, Seurat, or other publicly available packages and has been deposited at Github (bit.ly/4uIbvii).

